# Inferring hidden structure in multilayered neural circuits

**DOI:** 10.1101/120956

**Authors:** Niru Maheswaranathan, David B. Kastner, Stephen A. Baccus, Surya Ganguli

**Affiliations:** Neurosciences Graduate Program, Stanford University, Stanford, CA, USA; Department of Psychiatry, UCSF, San Francisco, CA, USA; Dept. of Neurobiology, Stanford University, Stanford, CA, USA; Dept. of Applied Physics, Stanford University, Stanford, CA, USA

## Abstract

A central challenge in sensory neuroscience involves understanding how neural circuits shape computations across cascaded cell layers. Here we develop a computational framework to reconstruct the response properties of experimentally unobserved neurons in the interior of a multilayered neural circuit. We combine non-smooth regularization with proximal consensus algorithms to overcome difficulties in fitting such models that arise from the high dimensionality of their parameter space. Our methods are statistically and computationally efficient, enabling us to rapidly learn hierarchical non-linear models as well as efficiently compute widely used descriptive statistics such as the spike triggered average (STA) and covariance (STC) for high dimensional stimuli. For example, with our regularization framework, we can learn the STA and STC using 5 and 10 minutes of data, respectively, at a level of accuracy that otherwise requires 40 minutes of data without regularization. We apply our framework to retinal ganglion cell processing, learning cascaded linear-nonlinear (LN-LN) models of retinal circuitry, consisting of thousands of parameters, using 40 minutes of responses to white noise. Our models demonstrate a 53% improvement in predicting ganglion cell spikes over classical linear-nonlinear (LN) models. Internal nonlinear subunits of the model match properties of retinal bipolar cells in both receptive field structure and number. Subunits had consistently high thresholds, leading to sparse activity patterns in which only one subunit drives ganglion cell spiking at any time. From the model’s parameters, we predict that the removal of visual redundancies through stimulus decorrelation across space, a central tenet of efficient coding theory, originates primarily from bipolar cell synapses. Furthermore, the composite nonlinear computation performed by retinal circuitry corresponds to a boolean OR function applied to bipolar cell feature detectors. Our general computational framework may aid in extracting principles of nonlinear hierarchical sensory processing across diverse modalities from limited data.

**Author Summary:** Computation in neural circuits arises from the cascaded processing of inputs through multiple cell layers. Each of these cell layers performs operations such as filtering and thresholding in order to shape a circuit’s output. It remains a challenge to describe both the computations and the mechanisms that mediate them given limited data recorded from a neural circuit. A standard approach to describing circuit computation involves building quantitative encoding models that predict the circuit response given its input, but these often fail to map in an interpretable way onto mechanisms within the circuit. In this work, we build two layer linear-nonlinear cascade models (LN-LN) in order to describe how the retinal output is shaped by nonlinear mechanisms in the inner retina. We find that these LN-LN models, fit to ganglion cell recordings alone, identify filters and nonlinearities that are readily mapped onto individual circuit components inside the retina, namely bipolar cells and the bipolar-to-ganglion cell synaptic threshold. This work demonstrates how combining simple prior knowledge of circuit properties with partial experimental recordings of a neural circuit’s output can yield interpretable models of the entire circuit computation, including parts of the circuit that are hidden or not directly observed in neural recordings.

## Introduction

### Motivation

Computational models of neural responses to sensory stimuli have played a central role in addressing fundamental questions about the nervous system, including how sensory stimuli are encoded and represented, the mechanisms that generate such a neural code, and the theoretical principles governing both the sensory code and underlying mechanisms. These models often begin with a statistical description of the stimuli that precede a neural response such as the spike-triggered average (STA) [1, 2] or covariance (STC) [3–8]. These statistical measures characterize to some extent the set of effective stimuli that drive a response, but do not necessarily reveal how these statistical properties relate to cellular mechanisms or neural pathways. Going beyond descriptive statistics, an explicit representation of the neural code can be obtained by building a model to predict neural responses to sensory stimuli.

A classic approach involves a single stage of spatiotemporal filtering and a time-independent or static nonlinearity; these models include linear-nonlinear (LN) models with single or multiple pathways [1, 9–11] or generalized linear models (GLMs) with spike history feedback [12, 13]. However, these models do not directly map onto circuit anatomy and function. As a result, the interpretation of such phenomenological models, as well as how they precisely relate to underlying cellular mechanisms, remains unclear. Ideally, one would like to generate more biologically realistic models of sensory circuits, in which sub-components of the model map in a one-to-one fashion onto cellular components of neurobiological circuits [14]. For example, model components such as spatiotemporal filtering, thresholding, and summation are readily mapped onto photoreceptor or membrane voltage dynamics, synaptic and spiking thresholds, and dendritic pooling, respectively.

A critical aspect of sensory circuits is that they operate in a hierarchical fashion in which sensory signals propagate through multiple nonlinear cell layers [15–17]. Fitting models that capture this widespread structure using neural data recorded from one layer of a circuit in response to controlled stimuli raises significant statistical and computational challenges [18–22]. A key issue is the high dimensionality of both stimulus and parameter space, as well as the existence of hidden, unobserved neurons in intermediate cell layers. The high dimensionality of parameter space can necessitate prohibitively large amounts of data and computational time required to accurately fit the model. One approach to address these difficulties is to incorporate prior knowledge about the structure and components of circuits to constrain the model [11, 21, 23–25]. Although prior knowledge of the exact network architecture and sequence of nonlinear transformations would greatly constrain the number of possible circuit solutions, such prior knowledge is typically minimal for most neural circuits.

In this work, we develop a computational framework that addresses these challenges and use it to learn hierarchical nonlinear models of ganglion cells in the salamander retina. In particular, we focus on models with three cell layers connected by two stages of linear-nonlinear processing (LN-LN models). As described below, the cell layers of these models map in one-to-one fashion onto the three principal cell layers of the retina: photoreceptors, bipolar cells, and retinal ganglion cells. We demonstrate that these models are both a more accurate description of the retinal code, as well as more amenable to biophysical interpretation. In particular, we find a match between the properties of subunits in the intermediate, hidden layer of our models and the properties of bipolar cells in the retina. Further analysis of our learned models reveals novel insight into retinal function, namely that, (1) transmission between every subunit and ganglion cell pair is well described by a high threshold expansive nonlinearity, (2) bipolar cells are sparsely active, (3) visual inputs are most decorrelated at the subunit layer, pre-synaptic to ganglion cells, and (4) the composite computation performed by the retinal ganglion cell output corresponds to a boolean OR function of bipolar cell feature detectors. Collectively, these results shed light on the nature of hierarchical nonlinear computation in the retina. Our computational framework is general, however, and we hope it will aid in providing insights into hierarchical nonlinear computations across the nervous system.

## Background

The retina is a classic system for exploring the relationship between quantitative encoding models and measurements of neurobiological circuit properties [26, 27]. Signals in the retina flow from photoreceptors through populations of horizontal, bipolar, and amacrine cells before reaching the ganglion cell layer.

To characterize this complex multilayered circuitry, many studies utilize descriptive statistics such as the spike-triggered average, interpreted as the average feature encoded by a ganglion cell [1–3]. Responses are often then modeled using a linear-nonlinear (LN) framework (schematized in Figure 1a). A major reason for the widespread adoption of LN models is their high level of tractability; learning their parameters can be accomplished by solving a simple convex optimization problem [2], or alternatively, estimated using straightforward reverse correlation analyses [1]. However, LN models have two major drawbacks: it is difficult to map them onto biophysical mechanisms in retinal circuitry, and they do not accurately describe ganglion cell responses across diverse stimuli. Regarding mechanisms, the spatiotemporal linear filter of the LN model is typically interpreted as mapping onto the aggregate sequential mechanisms of phototransduction, signal filtering and transmission through bipolar and amacrine cell pathways, and summation at the ganglion cell, while the nonlinearity is mapped onto the spiking threshold of ganglion cells. Regarding accuracy, while previous studies have found that these simple models can, for some neurons, capture most of the variance of the responses to low-resolution spatiotemporal white noise [9, 12, 20], they do not describe responses to stimuli with more structure such as natural scenes [13, 28–30]. A likely reason for these drawbacks are the nonlinearities within the retina. There can be strong rectification of signals that occurs pre-synaptic to ganglion cells [15, 31–33], breaking the assumption of composite linearity in the pathway from photoreceptors just up to the ganglion cell spiking threshold [17]. Indeed, nonlinear spatial integration within ganglion cell receptive fields was first described in the cat retina [34] in *Y*-type ganglion cells. A hypothetical model for this computation was proposed as a cascade of two layers of linear-nonlinear operations (LN-LN) [35, 36]. If one keeps the mean luminance constant, avoiding light adaptation in photoreceptors, the first major nonlinearity is thought to lie at the presynaptic terminal of the bipolar to ganglion cell synapse. Ganglion cells pool over multiple bipolar cell inputs, each of which can be approximated as linear-nonlinear components, termed subunits of the ganglion cell^1^. The second LN layer corresponds to summation or pooling across multiple subunits at the ganglion cell soma, followed by a spiking threshold. The subunit nonlinearities in these models have been shown to underlie many retinal computations including latency encoding [27], object motion sensitivity [37], and sensitivity to fine spatial structure (such as edges) in natural scenes [32]. Figure 1 shows a schematic of the LN-LN cascade and its mapping onto retinal anatomy. Functionally, these models with multiple nonlinear pathways are both more amenable to interpretation and potentially provide a more accurate description of ganglion cell responses.

**Fig 1.**
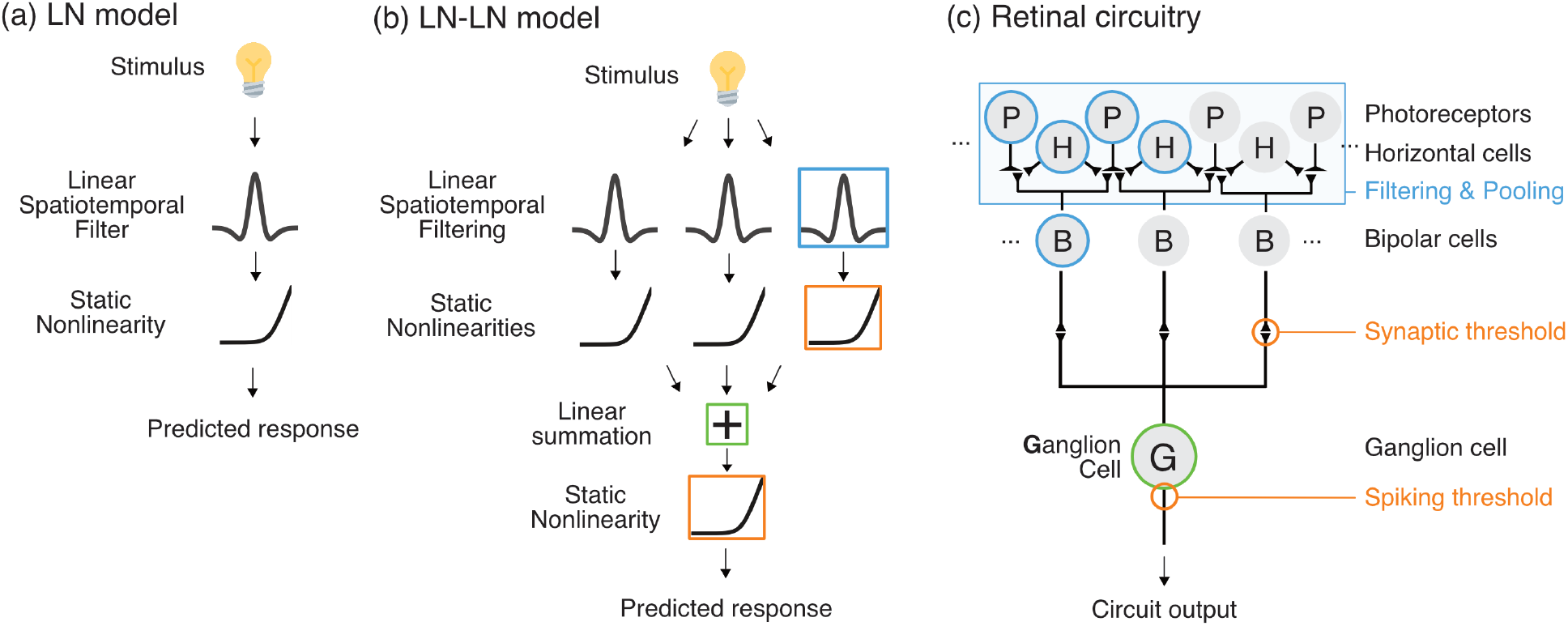
Schematics of the LN and LN-LN models and corresponding retinal circuitry, (a) The linear-nonlinear (LN) model consists of a single linear spatiotemporal filter followed by a static nonlinearity, (b) The LN-LN cascade contains a bank of LN subunits, whose outputs are pooled at a second linear stage before being passed through a final nonlinearity, (c) The LN-LN model mapped on to a retinal circuit. The first LN stage consists of bipolar cell subunits and the bipolar-to-ganglion cell synaptic threshold. The second LN stage is pooling at the ganglion cell, plus a spiking threshold.

### Related work

Early work on characterizing these multiple pathways motivated the use of the significant eigenvectors of the spike-triggered covariance (STC) matrix as the set of features that drives a cell, focusing on low-dimensional full field flicker stimuli [10, 38] to reduce the amount of data required for accurately estimating these eigenvectors. Significant STC eigenvectors will span the same linear subspace as the true biological filters that make up the pathways feeding onto a ganglion cell [3, 39–41]. However, the precise relationship between these eigenvectors (which obey a biologically implausible orthogonality constraint) and the individual spatiotemporal filtering properties intrinsic to multiple parallel pathways in a neural circuit remains unclear.

Instead, we take the approach of directly fitting a hierarchical, nonlinear, neural model, enabling us to jointly learn a set of non-orthogonal, biophysically plausible set of pathway filters, as well as an arbitrary, flexible nonlinearity for each pathway. Much recent and complementary work on fitting such models used simplifying assumptions in order to make model fitting tractable. For example, assuming the subunits are shifted copies of a template results in models with a single convolutional subunit filter [21, 23, 24]. However, this obscures individual variability in the spatiotemporal filters of subunits of the same type across visual space, which has been shown to be functionally important in increasing retinal resolution [42]. Restricting the number of subunits to a small number, such as two, yields models with separate ON-and OFF-pathways [18] but these do not meaningfully map onto anatomical pathways. Another common assumption is that the subunit nonlinearities have a particular form, such as quadratic [11, 25] or sigmoidal [43]. Fitting multi-layered models with convolutional filters and fixed nonlinearities has also been successfully used to describe retinal responses to natural scenes [30], although this work maximizes predictive accuracy at the expense of a one-to-one mapping of model components onto retinal circuit elements. Finally, other work focuses on particular ganglion cell types with a small number of inputs [22], constrains the input stimulus to a low-dimensional subspace (such as two halves of the receptive field [44]), or constrains the coefficients of receptive fields to be non-negative [45], thus discarding known properties of the inhibitory surround.

In this work, we do not make assumptions about or place restrictions on the number or tiling of subunit filters, the shapes of the subunit nonlinearities, the sign of receptive field elements, or the stimulus dimensionality. Instead, we place penalties on model parameters that encourage subunit filters to conform more closely to known statistical properties, namely that they should be sparse (contain few non-zero elements) and low-rank (approximately spatiotemporally separable) [46, 47], properties that are common to receptive fields in a wide variety of sensory systems. We describe computational methods based on proximal consensus algorithms, described below, that allow us to utilize this prior knowledge about model parameters in a computationally and statistically efficient manner to perform both spike-triggered analyses and fit hierarchical nonlinear models using much less data than otherwise required.

## Results

### Regularization techniques for model fitting

Fitting encoding models or computing descriptive statistics requires collecting enough neural data to constrain model parameters. Limited recording time, higher resolution stimuli, and more varied experimental conditions all necessitate being able to do more with less data. Depending on these factors, collecting enough data for fitting LN models is a challenge, and for an LN-LN model the problem is even more extreme. Regularization is also an issue for estimating descriptive statistics, like the spike-triggered average (STA) and covariance (STC), especially when the dimensionality of the stimulus is high.

To better constrain parameters of sensory encoding models as well as descriptive statistics we incorporate prior knowledge about simple statistical properties of the parameters in the form of regularization penalties. We encourage spatiotemporal filters in LN or LN-LN models, or the STA or STC eigenvectors to be sparse and approximately space-time separable (low-rank) by applying *ℓ*_1_-norm and nuclear norm penalties, respectively. These penalties have the additional benefit that they can efficiently be incorporated into optimization procedures using proximal algorithms (see Methods for details). Proximal algorithms [48, 49] are an appropriate choice for this problem because they can flexibly incorporate different penalty terms and they efficiently scale as the amount of data or stimulus dimensionality increases. In particular, they are better suited for problems with non-smooth terms (such as the regularization penalties we use) compared to gradient descent [48]. As a proof of concept for using these specific penalties, we utilized them to regularize the spike-triggered average (STA) (Eq. 1 in Methods) or eigenvectors of the spike-triggered covariance (STC) matrix (Eq. 3 in Methods) for retinal data.

Figure 2a compares a regularized spike-triggered average with the raw, un-regularized STA. For long recordings, the regularized STA closely matches the raw STA, while for short recordings the regularized STA has less high frequency noise and retains much of the structure observed if the STA had been estimated using more data. Figure 2b shows the performance of the regularized STA across different regularization weights, scanned over a broad range, demonstrating that performance is largely insensitive to the strengths of the weights of the *ℓ*_1_ and nuclear norm penalty functions. Thus regularization weights need not be fine tuned to achieve superior performance. We further quantified the performance of the regularized STA by employing it as the linear filter of an LN model, and found that with regularization, about 5 minutes of recording was sufficient to achieve the performance obtained through 40 minutes of recording without regularization (Figure 2c).

**Fig 2.**
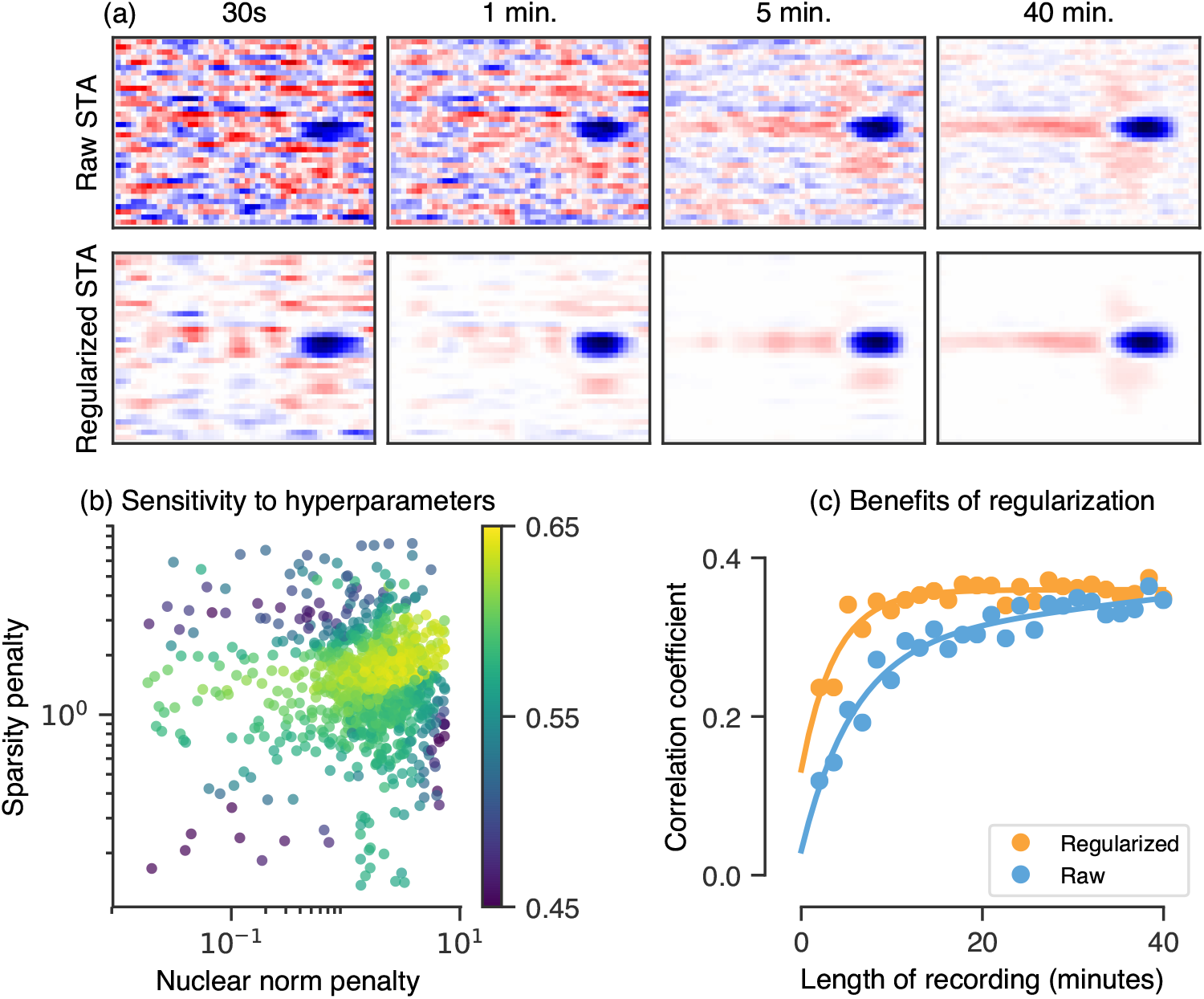
Regularization for estimating receptive fields (via a regularized spike-triggered-average). (a) Top row: the raw spike-triggered average computed using different amounts of data (from left to right, 30s to 4Omin), bottom row: the regularized spike-triggered average computed using the same amount of data as the corresponding column, (b) Performance (log-likelihood) as a function of two regularization weights, the nuclear norm (x-axis, encourages low-rank structure) and the *ℓ*_1_-norm (y-axis, encourages sparsity), (c) Correlation coefficient between the firing rate of a retinal ganglion cell and LN model whose filter is fixed to be a regularized or raw (un-regularized) STA, as a function of the amount of training data for estimating the STA (length of recording).

Beyond the STA, we also extended our framework to regularize essential information content in the eigenvectors of the sample STC matrix. Figure 3 demonstrates the improvement in our ability to estimate the relevant subspace spanned by significant STC eigenvectors, both in terms of the qualitative improvement in eigenvectors for an example cell (Fig. 3a) and quantified across the population (Fig. 3b). In Fig. 3a, we show the top regularized STC eigenvectors for different values of the nuclear norm (γ_*_) and *ℓ*_1_-norm (γ_1_) regularization penalties (Eq. 3 in Methods and Table S4). We score the performance of the STC subspace in Fig 3b in terms of how well stimuli, after projection onto the subspace, can be used to predict spikes, by computing the subspace overlap (defined in Methods) between the raw or regularized STC subspace and the best fit LN-LN subspace, obtained in the next section. This quantity ranges between zero for orthogonal subspaces and one for overlapping subspaces. Since the LN-LN subspace is the best subspace found by the LN-LN model for predicting spiking, a large subspace overlap between the regularized STC and LN-LN subspaces indicates the ability of regularized STC to find stimulus subspaces predictive of neural firing without actually fitting a model of the neuron.

**Fig 3.**
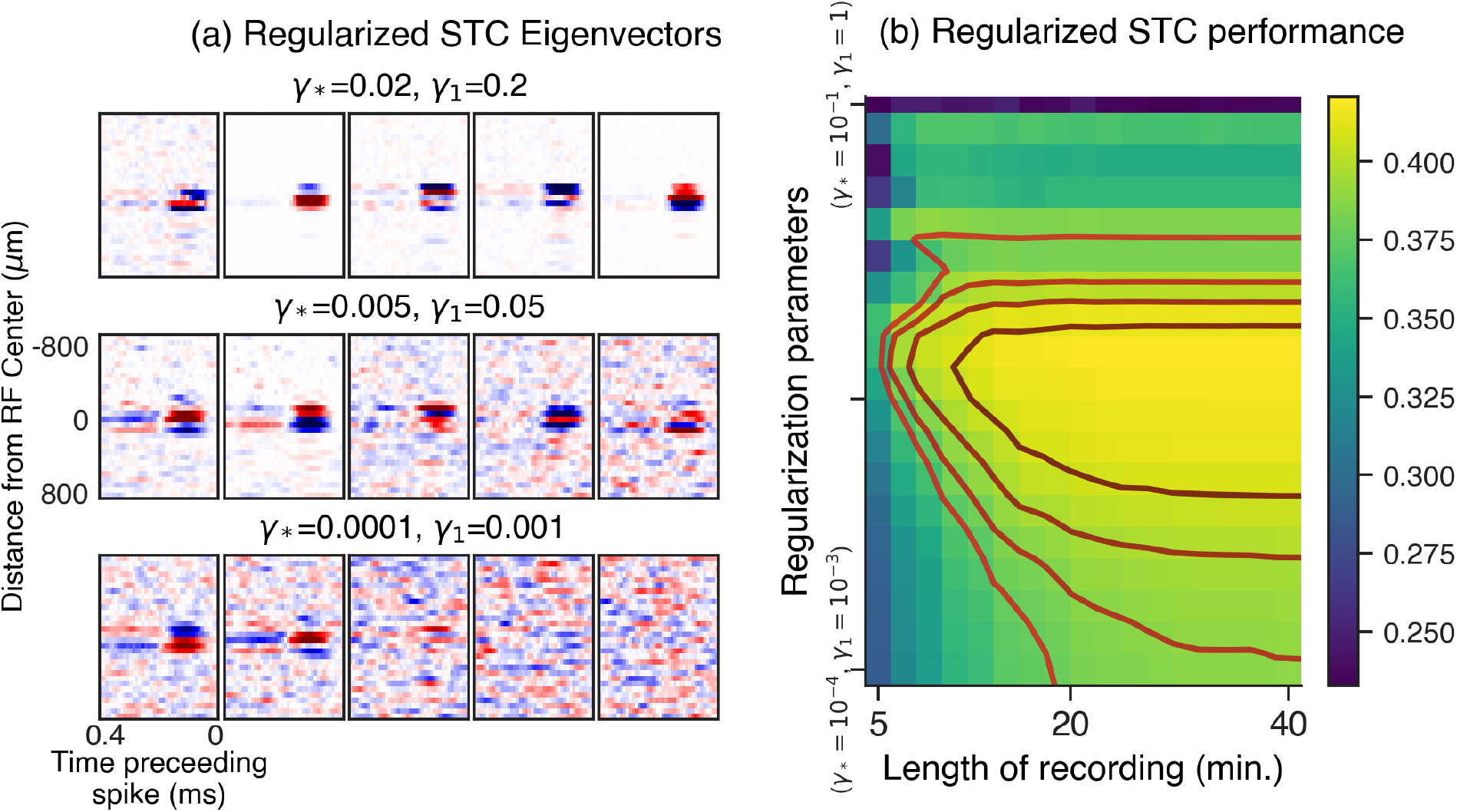
(a) Example panels of the output of our regularized spike-triggered covariance algorithm. Each panel contains the five most significant regularized eigenvectors of the STC matrix, reshaped as spatiotemporal filters. The bottom panel shows the result with no regularization added, and the upper panels show the result with increasing weights on the regularization penalties. Here *γ*_1_ is the regularization weight applied to an *γ*_1_ penalty encouraging sparsity, and *γ*_*_ is a regularization weight applied to a nuclear norm penalty, encouraging approximate spatiotemporal separability of the eigenvectors, when reshaped as spatiotemporal filters, (b) Summary across a population of cells. The heatmap shows the performance of regularized STC (measured as the subspace overlap with the best fit LN-LN subspace, see text for details). The y-axis in (b) represents a line spanning 3 orders of magnitude in two-dimensional regularization parameter space (Eq. 3 in Methods) (*γ*_*_,*γ*_1_), ranging from the point (*γ*_*_ = 10^−4^, *γ*_1_ = 10^−3^) to (*γ*_*_ = 10^−1^, *γ*_1_ = 1).

With appropriate regularization, one can recover the best predictive subspace using about 10 minutes of data; without regularization, one requires 40 minutes of data to recover a subspace with comparative predictive accuracy. Note that even for the full length of this experiment (40 minutes), regularization still improves our regularized STC estimate. Our much improved performance of regularized STC with limited amounts of data highlights the power of regularized eigenvector recovery using *γ*_1_ and nuclear norm penalties on the STC eigenvectors to find stimulus subspaces that are highly predictive of neural spiking, without ever directly fitting a parameterized model of the neuron.

Related work on regularizing the STA using Bayesian methods [46, 50, 51], often requires inversion of an *N*-by-*N* prior covariance matrix, where *N* is the dimension of stimulus space (a costly *O*(*N*^3^) operation) prohibitive for high-dimensional stimuli. For regularized STC, they would require the inversion of *N*^2^-by-*N*^2^ matrices, thus demanding a step of computational complexity *O*(*N*^6^) in each iteration of the internal loop required to learn the prior over STC matrices. In contrast, methods based on proximal consensus algorithms for regularizing STA or eigenvectors of STC matrices are much more efficient. The most costly step in terms of computational time for such regularization involves computing the proximal operator for the nuclear norm applied to each of the *d* columns of the matrix **X** in (3), when each column is viewed as an *N_s_* by *N_t_* spatiotemporal matrix. The proximal operator for the nuclear norm involves computing a singular value decomposition (SVD) of this matrix, and the computational cost of computing any such SVD is *O*([min(*N_s_*, *N_t_*)} × [max(*N_s_*, *N_t_*)]^2^). If the number of spatial bins *N_s_* and number of temporal bins *N_t_* are both of the same order of magnitude (and therefore each is of order 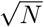), then the computational cost of a single SVD is *O*(*N*^3^/^2^). Thus the computational cost of STC eigenvector regularization through nuclear norm scales with stimulus dimension *N* as *N*^3/2^ in contrast to *N*^6^, as would be the case for empirical Bayes methods applied to STC matrices. One disadvantage of the regularization methods we use is that they cannot handle some of the more flexible priors employed in empirical Bayes methods.

### Learning hierarchical nonlinear models of the retinal response

Given the power of our regularization framework to efficiently learn the STA and STC, demonstrated above, we now apply this framework to learn LN-LN models. In the LN-LN architecture schematized in Figure 1, the stimulus is passed through a set of LN subunits. Each subunit filter is a spatiotemporal stimulus filter, constrained to have unit norm. The subunit nonlinearity is parameterized using a set of basis functions (Gaussian bumps) that tile the input space [12, 20] (see Methods). This parameterization is flexible enough that we could in principle learn, for each individual subunit, any smooth nonlinearity that can be expressed as a linear combination of our basis functions. The second LN layer pools subunits through weighted summation, followed by a spiking nonlinearity that we model using a parameterized soft rectifying function *r(x*) = *g* log(l + *e^xℓθ^*). Here *g* is an overall gain, and *θ* is a threshold. The full set of parameters for the model consists of the spatiotemporal subunit filters, the subunit nonlinearity parameters, and the gain and threshold of the final nonlinearity.

#### Model fitting and performance

We fit LN and LN-LN models to salamander ganglion cells using the 40 minute recording described previously. For both LN and LN-LN models we learned the model parameters by optimizing the sum of the log-likelihood of recorded spikes under a Poisson noise model [12] and the regularization terms. We chose the weights of the *ℓ*_1_ and nuclear norm regularization penalties, both for the LN and LN-LN models, through cross-validation on a small subset of cells, and then held these weights constant across all cells. Our subsequent results indicate that we do not have to fine tune these hyperparameters on a cell by cell basis to achieve good predictive performance. Finally, because different cells may have different numbers of functional subunits, for the LN-LN models, we chose the optimal number of subunits on a cell-by-cell basis by maximizing performance on held-out data through cross-validation. No additional structure was imposed on the subunits such as spatial repetition, overlap, or non-negativity.

We find that the LN-LN model significantly outperforms the LN model at describing responses of ganglion cells, for all recorded cells. Figure 4a shows firing rate traces for an example cell, comparing the recorded response (gray) with an LN model (blue) and an LN-LN model (red). We quantify the similarity between predicted and recorded firing rate traces using either the Pearson correlation coefficient or the log-likelihood of held-out data under the model. All log-likelihood values are reported as an increase over the log-likelihood of a fixed mean firing rate model, scaled by the firing rate (yielding units of bits/spike). Summarized across *n* = 23 recorded ganglion cells, we find that the LN-LN model yields a consistent improvement over the LN model using either metric (Fig. 4bc). Overall, this demonstrated performance improvement indicates that nonlinear spatial integration is fundamental in driving ganglion cell responses, even to white noise stimuli, and that an LN model is not sufficient to capture the response to spatiotemporal white noise. This salient, intermediate rectification that we identify computationally is consistent with previous measurements of bipolar-to-ganglion cell transmission in the retina [44, 52].

**Fig 4.**
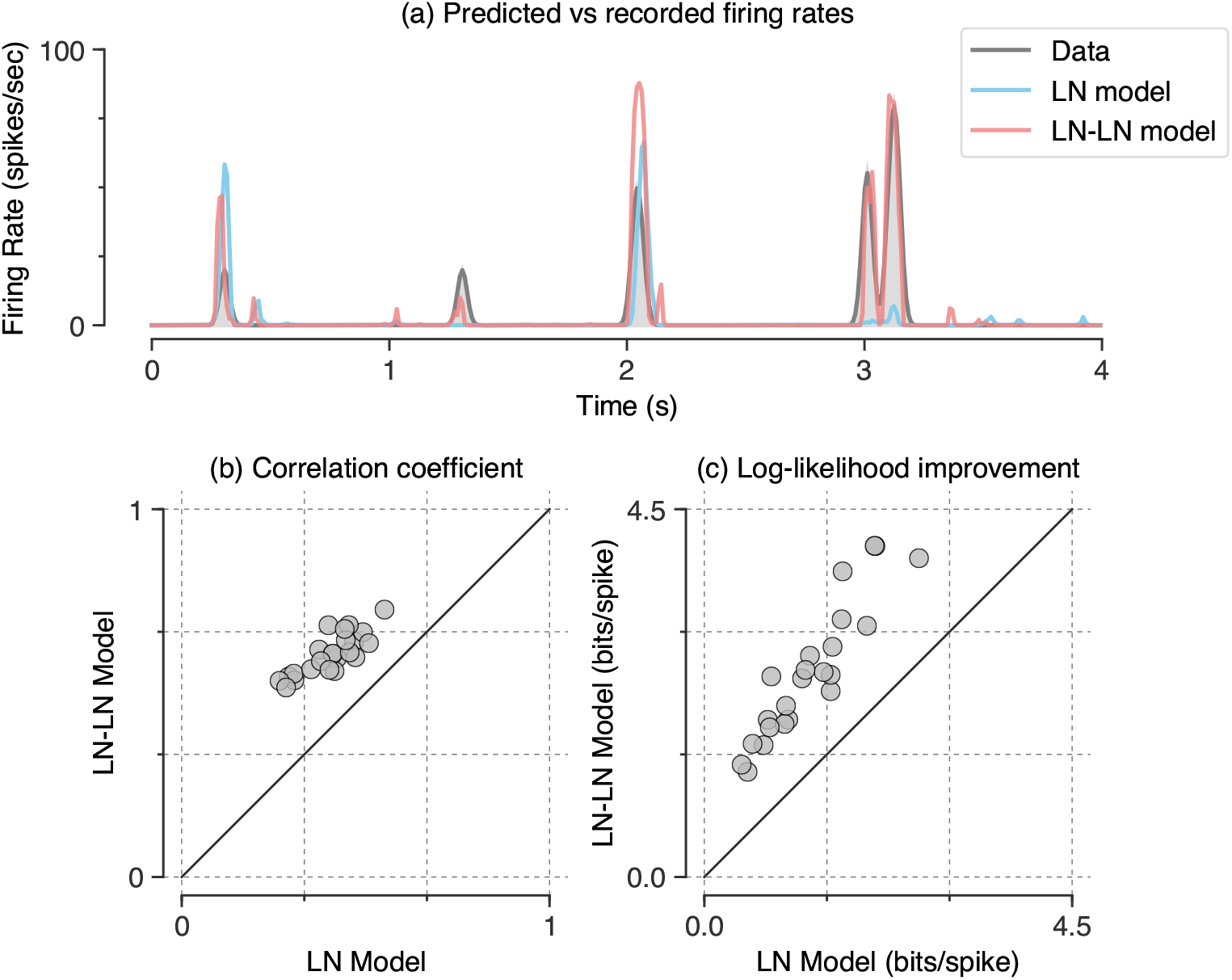
LN-LN models predict held-out response data better than LN models. (a) Firing rates for an example neuron. The recorded firing rate (shaded, gray), is shown along with the LN model prediction (dashed, green) and the LN-LN prediction (solid, red). (b) LN-LN performance on held out data vs. the LN model, measured using correlation coefficient between the model and held out data. Note that all cells are above the diagonal. (c) Same as in (b), but with the performance metric of log-likelihood improvement over the mean rate in bits per spike.

#### The internal structure of the learned models

Given the improved performance of our hierarchical nonlinear subunit models, we examined the internal structure of the models to assess their potential to reveal insights into retinal structure and computation. Figure 5 shows a visualization of the parameters learned for an example cell. Figure 5ab shows the parameters for the classical LN model, for comparison, while Figure 5cd shows the corresponding subunit filters and subunit nonlinearities in the first stage of the LN-LN model, fit to the same cell. The subunit filters had a similar temporal structure, but smaller spatial profiles compared to that of the LN model and the nonlinearities associated with each subunit were roughly monotonic with high thresholds (quantified below). These qualitative properties were consistent across all cells.

**Fig 5.**
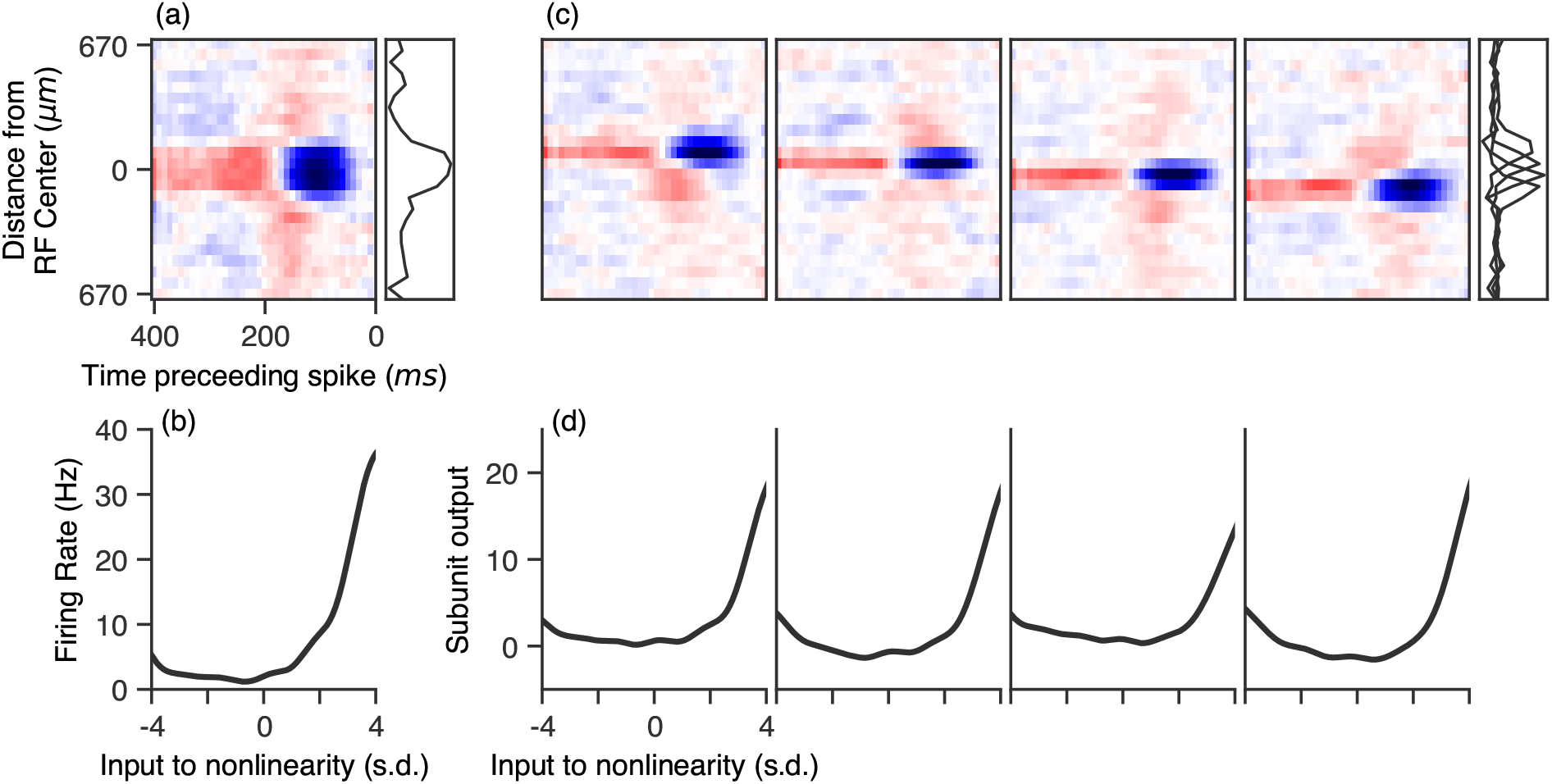
Example LN-LN model parameters fit to a recording of an OFF retinal ganglion cell. (a and b): LN-model parameters, consisting of a single spatial filter (a) and nonlinearity (b). (c and d) LN-LN model parameters. (c) First layer filters (top) and nonlinearities (bottom) of an LN-LN model fit to the same cell. Spatial profiles of filters are shown in gray to the right of the filters. The subunit filters have a much smaller spatial extent compared to the LN filter, but similar temporal profiles.

### Physiological properties of learned LN-LN models

Here we examine, in more detail, quantitative properties of learned LN-LN models that can be compared to physiological properties of the retina. In the next three sub-sections we find that model subunits quantitatively resemble bipolar cells in terms of receptive field properties and number, and that these intermediate subunits consistently have high-threshold nonlinearities.

#### Inferred hidden units quantitatively resemble bipolar cell receptive fields

Mapping the LN-LN model onto retinal anatomy leads us to believe that the first layer filters (some examples of which are shown in Figure 5c) should mimic or capture filtering properties pre-synaptic to bipolar cells in the inner retina. To examine this possibility, we compared the first layer model filters to properties of bipolar cell receptive fields. An example learned model subunit receptive field is shown in Fig. 6a, while a bipolar cell receptive field, obtained from direct intracellular recording of a bipolar cell, is shown in Figure 6b. Qualitatively, we found that the filters in the LN-LN model matched these bipolar cell RFs, as well as previously reported bipolar cell receptive fields [52, 53]: both had center-surround receptive fields with similar spatial extents. We further quantified the degree of space-time separability of the filters using the numerical or stable rank [54], which is a measure of rank insensitive to small amounts of noise in the matrix (a stable rank of one indicates the filter is exactly space-time separable). We found that the degree of space-time separability of recorded bipolar cell receptive fields and inferred model subunits were quite similar (1.28 ± 0. 01 and 1.39 ± 0.03, respectively), indicating that the nuclear norm penalty was not artificially reducing the rank of our model filters. This also demonstrates the advantage of using a soft rank penalty, such as the nuclear norm, as opposed to explicitly constraining the rank to any specific integer.

**Fig 6.**
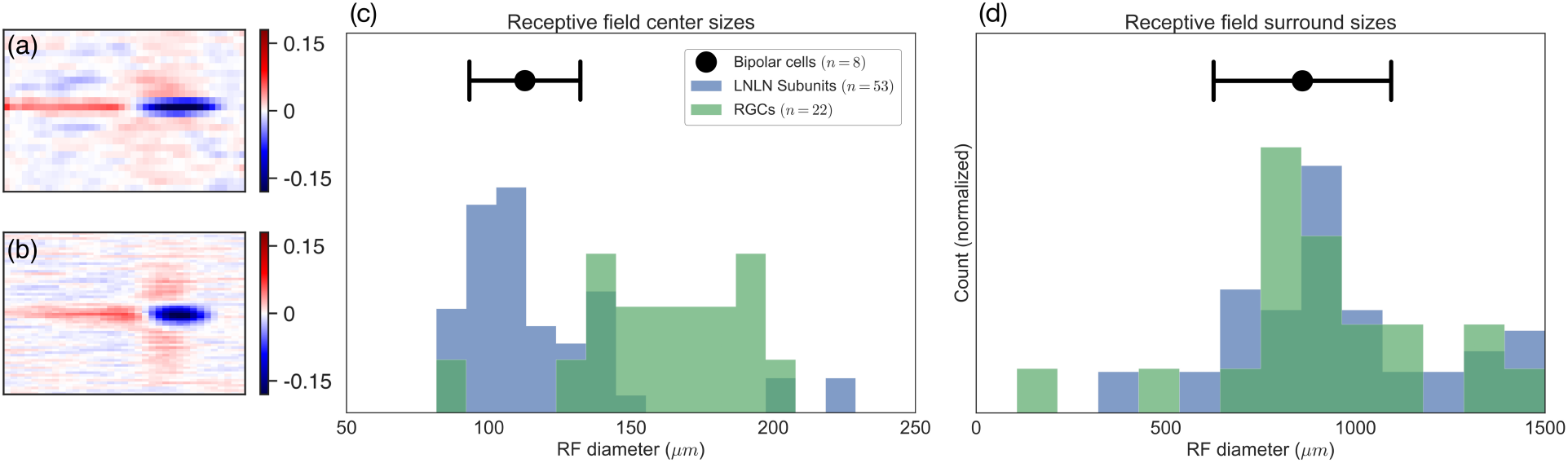
Comparison of subunit filter parameters with intracellular bipolar cell recordings. (a) An example subunit bipolar cell. (b) A recorded bipolar cell receptive field. (c) Receptive field centers sizes for subunit filters (blue), LN model filters (green), and recorded bipolar cells (black point). (c) Same as in (b), but with receptive field surround sizes.

To quantitatively compare these model-derived and ground-truth bipolar cell receptive fields, we fit the spatial receptive field with a difference of Gaussians function to estimate the RF center and surround sizes. We find the RF centers for the LN-LN subunit filters are much smaller than the corresponding LN model filter. Furthermore, the size of these LN-LN subunit centers matched the size of the RF center measured from intracellular recordings of *n* = 8 bipolar cells (Figure 6c). The recorded bipolar cells, LN model filters (ganglion cells), and LN-LN subunit filters all had similar surround sizes (Figure 6d). More example bipolar cell receptive fields are provided in Figure S2.

Note that this match between LN-LN model subunits and the RF properties of bipolar cells was not a pre-specified constraint placed on our model, but instead arose as an emergent property of predicting ganglion cell responses to white noise stimuli. These results indicate that our modeling framework not only enables higher performing predictive models of the retinal response, but can also reconstruct important aspects of the unobserved interior of the retina.

#### Number of inferred subunits

The number of subunits utilized in the LN-LN model for any individual cell was chosen to optimize model predictive performance on a held-out data set via cross-validation. That is, we fit models with different numbers of subunits and selected the one with the best performance on a validation set. We find that for models with more subunits than necessary, extra subunits are ignored (the learned nonlinearity for these subunits is flat, thus they do not modulate the firing rate).

Figure 7a shows the model performance, quantified as the difference between the LN-LN model and the LN model, across a population of cells as a function of the number of subunits included in the LN-LN model. We find that models with four to six subunits maximized model performance on held-out data. Note that the stimuli used here are one-dimensional spatiotemporal bars that have constant luminance across one spatial dimension. Thus each model subunit likely corresponds to the combination of multiple bipolar cell inputs whose receptive fields overlap a particular bar in the stimulus. Previous anatomical studies of bipolar cell density and axonal branching width [55, 56] as well as functional studies [45] suggest that a typical ganglion cell in the salamander retina receives input from 10–50 bipolar cells whose receptive fields are tiled across two dimensional space. The number of independently activated groups of such a two dimensional array of bipolar cells, in response to a one dimensional bar stimulus is then expected to be reduced from the total number of bipolar cells, roughly, by a square root factor: i.e. 25 → 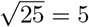. This estimate is largely consistent with the typical number of subunits in Figure 7a, required to optimize model predictive performance. This estimate also suggests that the large majority of bipolar-to-ganglion cell synapses are rectifying (strongly nonlinear), as linear connections are not uniquely identifiable in an LN-LN cascade. Indeed, in the salamander retina, strong rectification appears to be the norm [44, 52].

**Fig 7.**
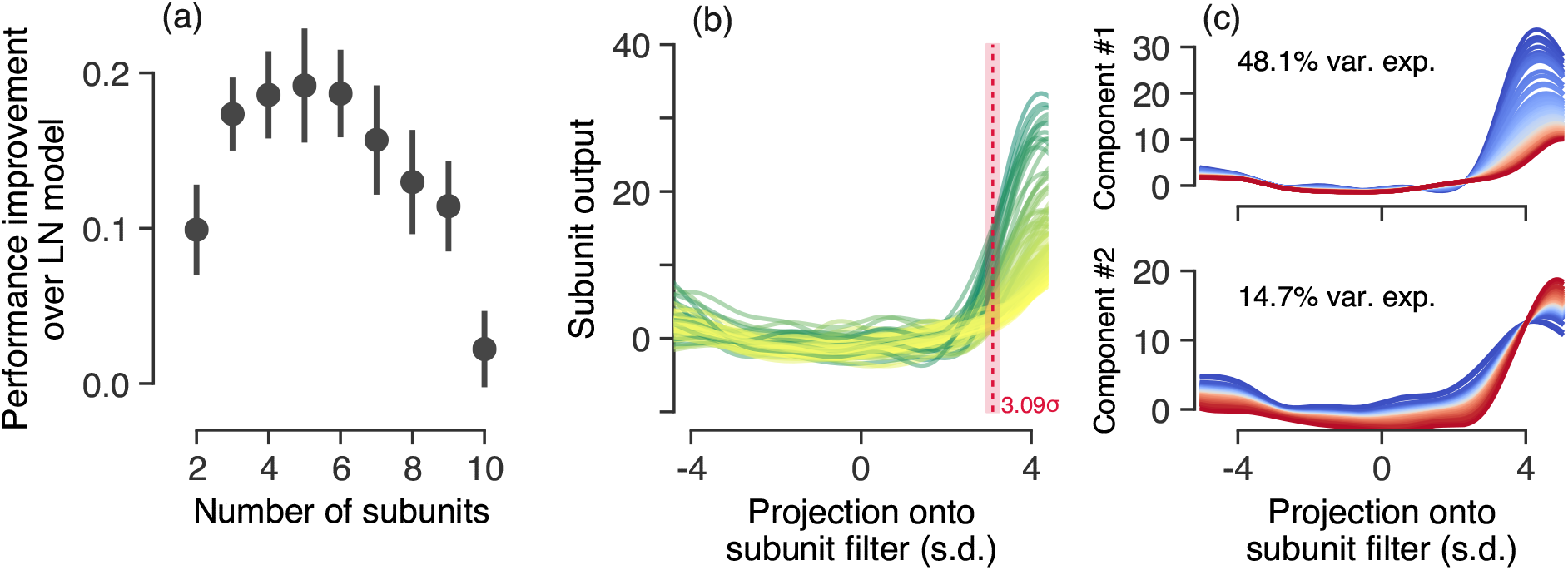
LN-LN model parameter analysis. (a) Performance improvement (increase in correlation coefficient relative to an LN model) as a function of the number of subunits used in the LN-LN model. Error bars indicate the standard error across 23 cells. (b) Subunit nonlinearities learned across all ganglion cells. For reference the white noise input to a subunit nonlinearity has standard deviation 1, which sets the scale of the x-axis. Red line and shaded fill indicate the mean and s.e.m. of nonlinearity thresholds (see text for details). (c) Visualization of the principal axes of variation in subunit nonlinearities by adding or subtracting principal components from the mean nonlinearity. (top) The principal axis of variation in subunit nonlinearities results in a gain change, while (bottom) the second principal axis corresponds to a threshold shift. These two dimensions captured 63% of the nonlinearity variability across cells.

#### LN-LN models have subunit nonlinearities with consistently high thresholds

The nonlinearities for all of the measured subunits are overlaid in Figure 7b. Each LN-LN subunit nonlinearity takes as input the projection of the stimulus onto the corresponding subunit spatiotemporal filter. Since the stimulus components are white noise with unit standard deviation, and the spatiotemporal filter is constrained to have unit norm, the projection of the stimulus onto the filter has a standard Normal distribution, thus we can compare nonlinearities on a common axis. Despite the fact that the model could separately learn an arbitrary function over the input for each subunit nonlinearity, we find that the nonlinearities are fairly consistent across the different subunits of many cells. Subunit nonlinearities look roughly like thresholding functions, relatively flat for most inputs but then increasing sharply after a threshold. We quantified the threshold as input for which the nonlinearity reaches 40% of the maximum output, across *n* = 92 model-identified subunits the mean threshold was 3.09 ± 0.14 (s.e.m.) standard deviations. We additionally computed LN thresholds for ganglion cells and found that they were similarly consistent across the population (3.22 ± 0.11 standard deviations). We decomposed the set of nonlinearities using principal components analysis and show the two primary axes of variation in Figure 7c. The primary axis of variation results in a gain change, while the secondary axis induces a threshold shift. Due to the high thresholds of these nonlinearities, subunits only impact ganglion cell firing probability for large input values. The slight rise on the left side of the nonlinearities is likely due to weak ON-inputs to the ganglion cell, and a stimulus ensemble that drives the ON-pathways more strongly [38, 57] may be necessary to uniquely identify them. Our ganglion cell population consisted of 18 Fast-Off, 4 Slow-Off, and 1 On cell (classification shown in Figure S3). We did not find significant differences across cell types in terms of the number of identified subunits or the subunit thresholds.

### Computational properties of learned LN-LN models

We now turn from a quantitative analysis of the physiological properties of the retina, described above, to their implications in terms of the computational function of the retina in processing visual stimuli. In particular, in the next two sub-sections we predict that the dominant contribution to stimulus decorrelation in efficient coding theory occurs at the bipolar cell synaptic threshold, and that the composite function computed by a retinal ganglion cell corresponds to a logical OR of its bipolar cell inputs.

#### Stimulus decorrelation at different stages in hierarchical retinal processing

Natural stimuli have highly redundant structure. Efficient coding theories [58, 59] state that sensory systems ought to remove these redundancies in order to efficiently encode natural stimuli. The simplest such redundancy is that nearby points in space and time contain similar, or correlated, luminance levels [60]. The transmission of such correlated structure would thus be highly inefficient. Efficient coding has been used to explain why responses are much less correlated than natural scenes, although the mechanistic underpinnings of decorrelation in the retina remain unclear.

Early work [61] suggested a simple mechanism: the linear center-surround receptive field of ganglion cells (and more recently, of bipolar cells [62]) could contribute to redundancy reduction simply by transmitting only differences in stimulus intensity across nearby positions in space. However, it was recently shown [63] using LN models that most of the decorrelation of naturalistic stimuli in the retina could be attributed to ganglion cell nonlinearities, as opposed to linear filtering. Given that we fit an entire layer of subunits pre-synaptic to each ganglion cell layer, we can analyze the spatial representation of naturalistic images at different stages of hierarchical retinal processing, thereby localizing the computation of decorrelation to a particular stage in the model.

To do so, we generated the response of the entire population of model subunits to a spatial stimulus similar to previous work [63], namely spatially pink noise, low pass filtered in time. We computed the correlation of stimulus intensities as a function of spatial distance, as well as the correlation between pairs of model units as a function of spatial distance. We examined pairs of units across different stages of the LN-LN model: after linear filtering by the subunits, after the subunit nonlinearity, and finally at the ganglion cell firing rates (the final stage). Figure 8 shows that the correlation at these stages drops off with distance between either the subunits or ganglion cells (with distance measured between receptive field centers). Similar to previous work [63], we find that most of the decorrelation is due to nonlinear processing, as opposed to linear filtering. However, our model predicts that this decorrelation is primarily due to the *subunit* nonlinearities, as opposed to ganglion cell spiking nonlinearities. In fact, the correlation between the ganglion cell model firing rates slightly increases after pooling across subunits. The most decorrelated representation occurs just after thresholding at the subunit layer. In this manner, our modeling framework suggests a more precisely localized mechanistic origin for a central tenet of efficient coding theory. Namely, our results predict that the removal of visual redundancies, through stimulus decorrelation across space, originates primarily from high-threshold nonlinearities associated with bipolar cell synapses.

**Fig 8.**
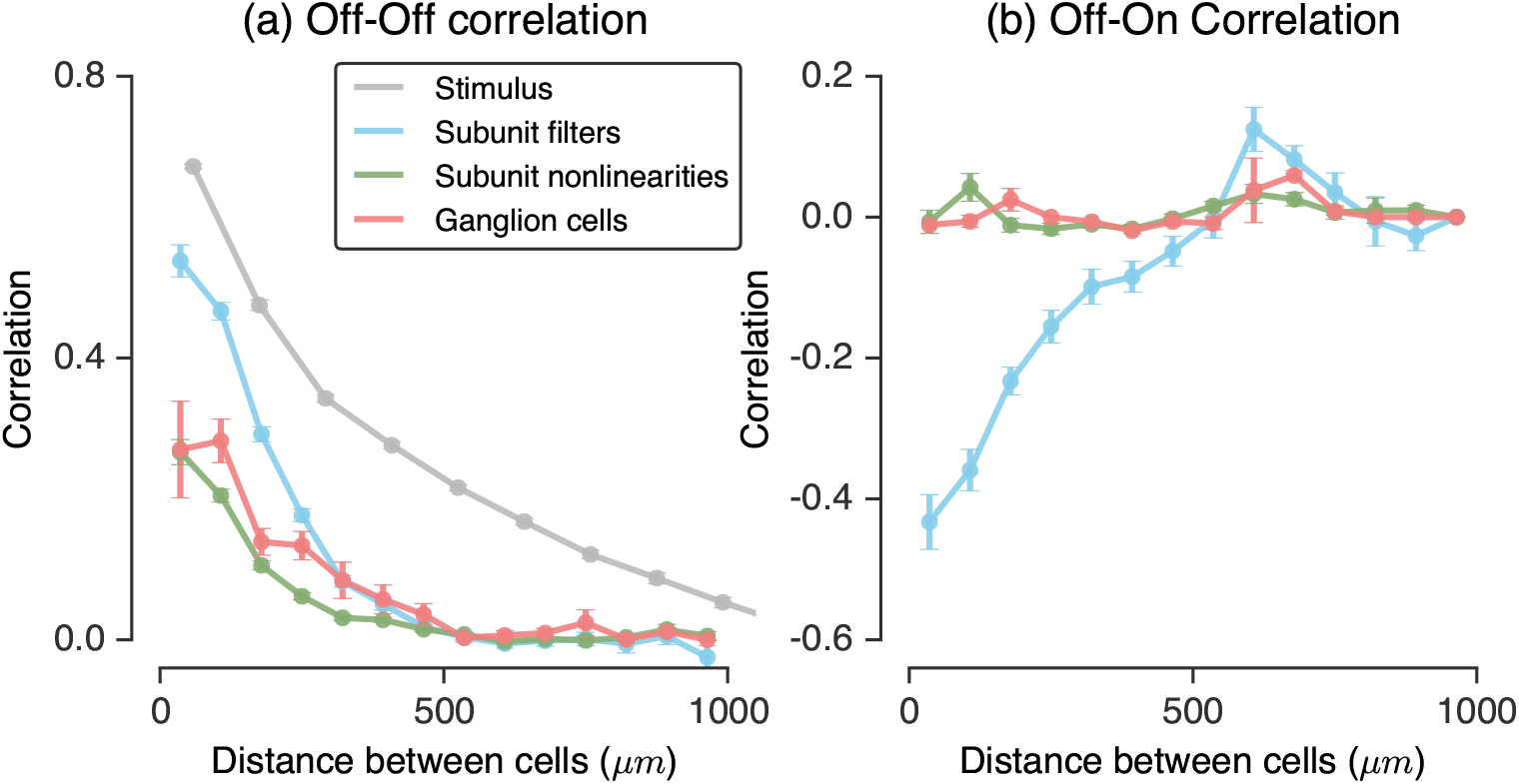
Decorrelation in LN-LN subunit models. A naturalistic (pink noise) stimulus was shown to a population of nonlinear subunits. The correlation in the population after filtering at the subunit layer (blue), after the subunit nonlinearity (green), and after pooling and thresholding at the ganglion cell layer (red), in addition to the stimulus correlations (gray) are shown. Left: the correlation as a function of distance on the retina for Off-Off cell pairs. Right: correlation for Off-On cell pairs. For each plot, distances were binned every 70μra, and error bars are the s.e.m. within each bin.

#### The nature of nonlinear spatial integration across retinal subunits

Retinal ganglion cells emit responses in sparse, temporally precise patterns [64], presumably to keep firing rates low thereby preserving energy [63, 65]. LN models can emulate sparse, precise firing in only one way: by using nonlinearities with high thresholds relative to the distribution of stimuli projected onto their linear filter. This way, only a small fraction of stimuli will cause the model to generate a response. Indeed, nonlinearities in LN models fit to ganglion cells have high thresholds [9]. LN-LN models, in contrast, can generate sparse responses using two qualitatively distinct operating regimes: either the subunit thresholds (first nonlinearity) could be high and the ganglion cell or spiking threshold (second nonlinearity) could be low, or the subunit thresholds could be low and spiking thresholds could be high. Both of these scenarios give rise to sparse firing at the ganglion cell output. However, they correspond to categorically distinct functional computations.

These various scenarios are diagrammed in Figure 9a-c. Each panel shows the response of a model in a two-dimensional space defined by the projection of the stimulus onto two subunit filters pre-synaptic to the ganglion cell (the two-dimensional space is easier for visualization, but the same picture holds for multiple subunits). We show the response as contours where the firing probability is constant (iso-response contours). Here, the subunit nonlinearities play a key role in shaping the geometry of the response contours, and therefore shape the computation performed by the cell. Note that the ganglion cell nonlinearity would act to rescale the output, but cannot change the shape of the contours. Therefore, it is fundamentally the subunit nonlinearities alone that determine the geometry of the response contours.

**Fig 9.**
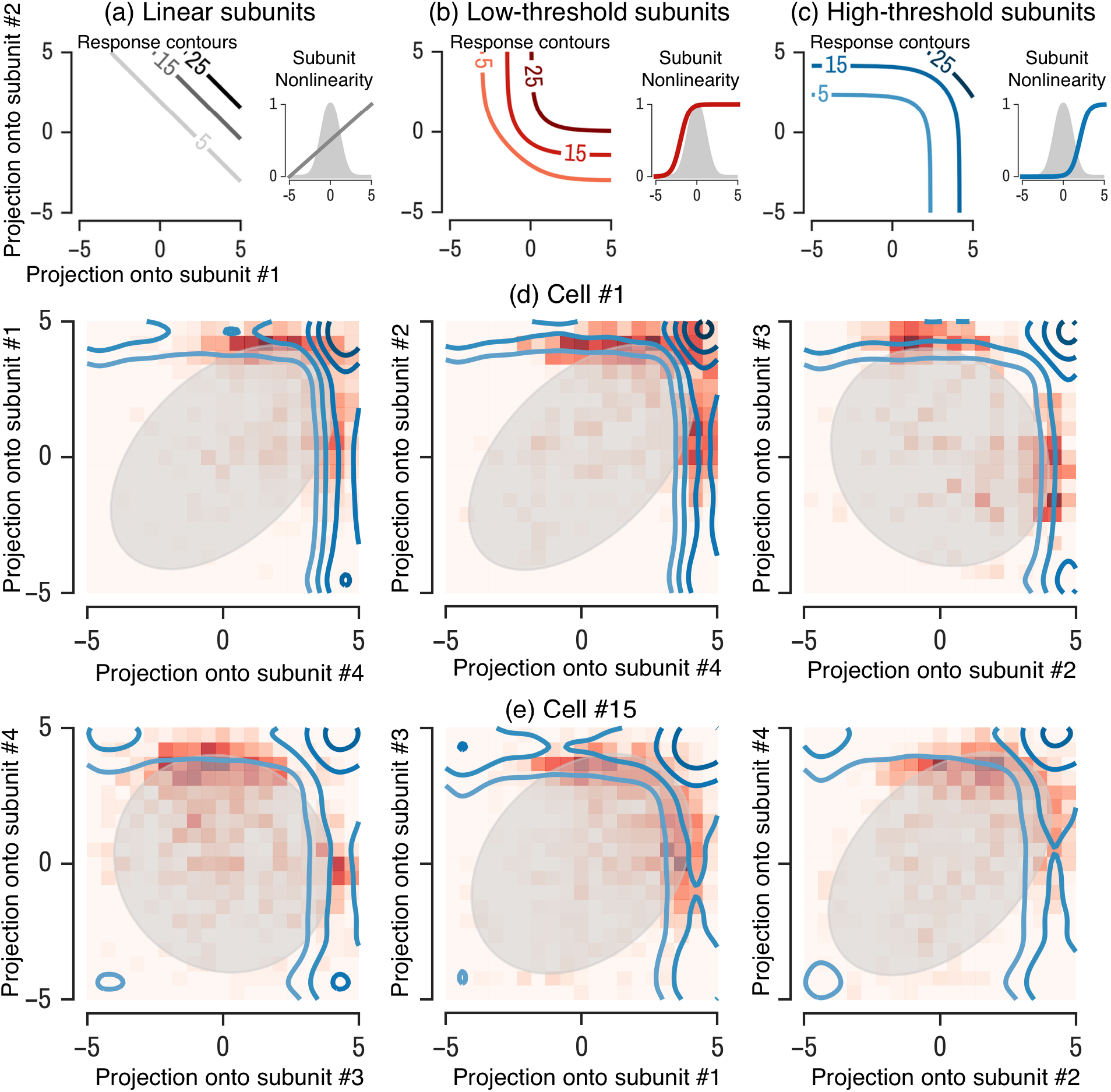
Visualization of subunit contours. Contours of equal firing probability are shown in a 2D space defined by the projection of the visual stimulus along each of two subunits. (a) Example contour plots for a model with low threshold subunit nonlinearities (inset) has concave contours. (b) A model with high threshold subunit nonlinearities has convex contours. (c & d) Contours from a model for two example ganglion cells, for three different pairs of subunits (left to right). In each panel, a histogram of the recorded firing rate is shown (red squares) as well as the stimulus distribution (gray oval).

Low-threshold subunit nonlinearities give rise to concave contours (Figure 9b), whereas high-threshold subunits give rise to convex contours (Figure 9c). Because final output rate is determined by the subunit and final thresholds, both of these descriptions could yield sparse firing output with the same overall rate (by adjusting the final threshold), but correspond to different computations. Low-threshold subunits can be simultaneously active across many stimuli, and thus yield spiking when subunits are simultaneously active (an AND-like combination of inputs). On the other hand, high-threshold subunits are rarely simultaneously active and thus usually only one subunit is active during a ganglion cell firing event, giving rise to an OR-like combination of inputs. By comparison, a cell that linearly integrates its inputs would have linear contours (Figure 9a).

In our models fit to retinal ganglion cells, we find all cells are much more consistent with the high threshold OR-like model. Subunit nonlinearities tend to have high thresholds, and therefore result in convex contours (shown for different pairs of subunits for two example cells in Figure 9d-e). For each example ganglion cell, we show the corresponding model contours along with the 2 standard deviation contour of the stimulus distribution (gray oval) and the empirical firing histogram (red checkers) in the 2D space defined by the projection of the stimulus onto a given pair of subunit filters identified by the LN-LN cascade model. Note that while the stimulus is uncorrelated (i.e. white, or circular), non-orthogonality of subunit filters themselves yield correlations in the subunit activations obtained by applying each subunit filter to the stimulus. Hence the stimulus distribution in the space of subunit activations (grey shaded ovals) is not circular. In all recorded cells, we find that the composite computation implemented by retinal ganglion cell circuitry corresponds to an OR function associated with high subunit thresholds (as schematized in Figure 9c). Moreover, both the AND computation and the linear model are qualitatively ruled out by the shape of the model response contours as well as the empirical firing histogram over subunit activations, which closely tracks the model response contours (i.e. the boundaries of the red histograms are well captured by the model contours).

These results are consistent with previous studies of nonlinear spatial integration in the retina. For example, Bollinger et. al. [44] discovered convex iso-response contours for a very simple two dimensional spatial stimulus, and Kaardal et. al. [43] performed an explicit hypothesis test between an AND-like and OR-like nonlinear integration over a low dimensional subspace obtained via the un-regularized STC eigenvectors, finding that OR outperformed AND. However, the techniques of [44] can only explore a low-dimensional stimulus space, whereas our methods enable the discovery of iso-response contours for high-dimensional stimuli. Moreover, in contrast to the hypothesis testing approach taken in [43], our general methods to learn LN-LN models reveal that an OR-model of nonlinear integration is a good model on an absolute scale of performance amongst all models in the LN-LN family, rather than simply being better than an AND-model.

#### A multi-dimensional view of cascaded retinal computation

A simple, qualitative schematic of the distinct computational regime in which retinal ganglion cells operate in response to white noise stimuli can be obtained by considering the geometry of the spike triggered ensemble in *N* dimensional space. In particular, the distribution of stimuli concentrates on a constant radius sphere in *N* dimensional stimulus space. More precisely, any high dimensional random stimulus realization **x** has approximately the same vector length, because the fluctuations in length across realizations, relative to the mean length, is 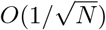. Thus we can think of all likely white-noise stimuli as occurring on the *N* – 1 dimensional surface of a sphere in *N* dimensional space. Each subunit filter can be thought of as a vector pointing in a particular direction in *N* dimensional stimulus space. The corresponding input to the subunit nonlinearity for any stimulus is the inner-product of the stimulus with the subunit filter, when both are viewed as *N* dimensional vectors. The high threshold of the subunit nonlinearity means that the subunit only responds to a small subset of stimuli on the sphere, corresponding to a small cap centered around the subunit filter. For a single subunit model (i.e. an LN model), the set of stimuli that elicit a spike then corresponds simply to this one cap (Figure 10a). In contrast, the OR like computation implemented by an LN-LN model with high subunit thresholds responds to stimuli in a region consisting of a union of small caps, one for each subunit (Figure 10b).

**Fig 10.**
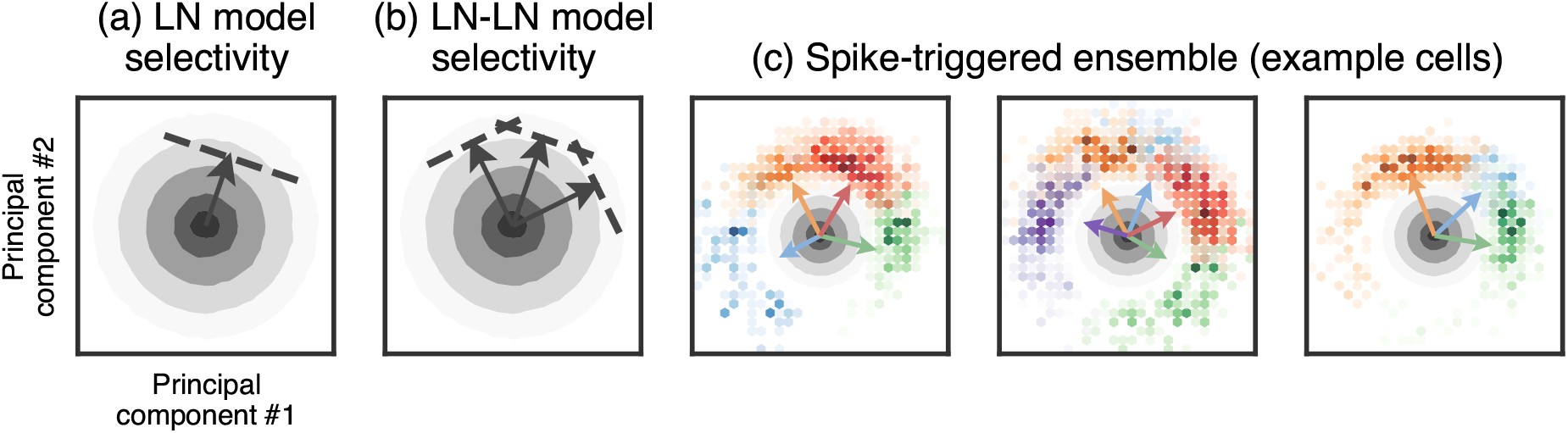
Stimulus selectivity in LN and LN-LN models. Each panel shows the raw stimulus distribution (gray contours) projected onto the top two principal components of the spike-triggered subunit activations (with subunits identified by the LN-LN model). The LN model (a) fires in response to stimuli in a single region, or cap, of stimulus space (indicated by the arrow and dashed threshold), whereas the LN-LN model (b) fires in response to a union of caps, each defined by an individual subunit, (c) Spike-triggered subunit activations for three representative cells are shown as colored histograms (colors indicate which model-identified subunit was maximally active during the spike), with the corresponding subunit filter directions shown as colored arrows (see text for details). Color intensity of the histogram indicates the probability density of the spike-triggered ensemble (STE), thus drops in intensity between changes in color indicate a multimodal STE, with high density modes centered near subunit filter directions.

To verify this conceptual picture, in Figure 10c we visualize a two-dimensional projection of the spike-triggered stimulus ensemble for three example ganglion cells, using principal components analysis of the spike-triggered subunit activations. That is, we project the spike-triggered ensemble onto the subunit filters identified in the LN-LN model, and subsequently project those subunit activations onto the two dimensions that capture the most variance in subunit activations. Note that this is different from just taking the top two principal components of the STC matrix, as the top STC component is typically the *average* of the subunit filters [10], which does not differentiate the subunit activations. This procedure identifies a subspace that captures the radial spread of subunit filters in high-dimensional stimulus space. We find that the spike-triggered ensemble projected onto this subspace (Figure 10c) curves around the radial shell defined by the stimulus distribution, and matches the conceptual picture shown in Figure 10b. For ease of visualization, we colored elements in the spike-triggered ensemble by which LN-LN model subunit was maximally active during that spike, and we normalize the spike-triggered histogram by the raw stimulus distribution (gray ovals). This picture provides a simple, compelling view for why LN models are insufficient to capture the retinal response to white noise, and further visualizes the aspect of retinal computation LN-LN models capture that the LN model does not: ganglion cells encode the union of different types of stimuli, with each stimulus type having large overlap with precisely one subunit filter.

## Discussion

In summary, we provide a computational framework to model stimulus driven neural processing in circuits with multiple parallel, hierarchical nonlinear pathways using limited experimental data. This framework elucidates relationships between biophysical circuit properties (spatiotemporal filtering properties of individual pathways, and nonlinear pooling of such pathways across multiple cell layers) and the statistical structure of the spike-triggered stimulus ensemble. We found that models employing two stages of linear and nonlinear computation, namely LN-LN models, demonstrated a robust improvement over the classical standard of LN models at predicting responses to white noise across a population of ganglion cells.

Beyond performance considerations alone, the gross architecture of the LN-LN model maps directly onto the hierarchical, cascaded, anatomy of the retina, thereby enabling the possibility that we can generate quantitative hypotheses about neural signal propagation and computation in the unobserved interior of the retina simply by examining the structure of our model’s interior. Since learning our model only requires measurements of the inputs and outputs to the retinal circuit, this approach is tantamount to the computational reconstruction of unobserved hidden layers of a neural circuit. The advantage of applying this method in the retina is that we can experimentally validate aspects of this computational reconstruction procedure.

Indeed, using intracellular recordings of bipolar cells, we found that our learned subunits matched properties of bipolar cells, both in terms of their receptive field center-surround structure, and in terms of the approximate number of bipolar cells connected to a ganglion-cell. However care must be taken not to *directly* identify the learned subunits in our model with bipolar cells in the retina. Instead, they should be thought of as *functional* subunits that reflect the combined contribution of not only bipolar cells, but also horizontal cells and amacrine cells that sculpt the composite response of retinal ganglion cells to stimuli. Nevertheless, the correspondence between subunits and bipolar cell RFs (which are also shaped by horizontal cells), suggests learning functional subunits that loosely correspond to the composite effect bipolar cells and associated circuitry have on ganglion-cell synapses is important in explaining the overall ganglion cell response, even to white-noise stimuli.

The interior of our models also reveal several functional principles underlying retinal processing. First, all subunits across all cells had strikingly consistent nonlinearities corresponding to monotonically increasing threshold-like functions with very high thresholds. This inferred biophysical property yields several important consequences for neural signal processing in the inner retina. First, it predicts that subunit activation patterns are sparse across the ensemble of stimuli, with typically only one subunit actively contributing to any given ganglion cell spike. Second, it predicts that the dominant source of stimulus decorrelation, a central tenet of efficient coding theory, has its mechanistic origin at the first strongly nonlinear processing stage of the retina, namely in the synapse from bipolar cells to ganglion cells. Third, it implies that the composite function computed by individual retinal ganglion cells corresponds to a Boolean OR function of bipolar cell feature detectors.

Taken together, our framework provides a unified way to estimate hierarchical nonlinear models of sensory processing by combating both the statistical and computational curses of dimensionality associated with learning such models. When applied to the retina, these techniques demonstrably recover known properties in the interior of the retina without requiring direct measurements of these properties. Moreover, by identifying candidate mechanisms for cascaded nonlinear computation in retinal circuitry, our results provide a higher resolution view of retinal processing compared to classic LN models, thereby setting the stage for the next generation of efficient coding theories that may provide a normative explanation for such processing. For example, considerations of efficient coding have been employed to explain aspects of the linear filter [66] and nonlinearity [63] of retinal ganglion cells when viewed through the coarse lens of an LN model. An important direction for future research would be the extension of these basic theories to more sophisticated ones that can explain the higher resolution view of retinal processing uncovered by our learned LN-LN models. Principles that underlie such theories of LN-LN processing might include subthreshold noise rejection [67, 68], sensitivity to higher order statistical structure in natural scenes, and energy efficiency [65]. Indeed the ability to extract these models from data in both a statistically and computationally efficient manner constitutes an important step in the genesis and validation of such a theory.

Another phenomenon robustly observed in the retina is adaptation to the luminance and contrast of the visual scene. Adaptation is thought to be a critical component of the retinal response to natural scenes [28], and a promising direction for extensions of our work would be to include luminance and contrast adaptation in subunit models. Luminance adaptation (adapting to the mean light intensity) is mediated by photoreceptor cells, and could be modeled by prepending a simple photoreceptor model (e.g. [69]) to an LN-LN model. There are two major sites of contrast adaptation, at the bipolar-to-ganglion cell synapse [70, 71] and at the spiking mechanism of ganglion cells [70, 72]. Extending the simple thresholding nonlinearities in our model with a dynamical model of adaptation (e. g. [19]) is a first step towards understanding the interaction between nonlinear subunits and adaptation.

While our work utilized white noise stimuli, our methods do not make assumptions about the stimulus statistics and will likely generalize to other stimulus distributions. In particular, stimuli that differentially activate subunits will be the most effective at differentiating LN and LN-LN models. Stimuli with coarse spatial resolution will not differentially activate subunits within the receptive field, thus are a poor choice for studying nonlinear spatial integration. However, fine textures as present in natural stimuli, are very likely to activate these nonlinear mechanisms in the retina, and thus are a critical component for understanding vision in the context of ethologically relevant stimuli.

The computational motifs identified by LN-LN models are likely to generalize across different species because they rely on a few key properties. For example, our predictions about the primary source of decorrelation in the retina rely on three features of the underlying circuitry identified by LN-LN models: (a) bipolar cell receptive fields are smaller than those of ganglion cells, (b) bipolar cell receptive field centers are largely non-overlapping, and (c) bipolar cell synapses have high thresholds. In addition, the logical OR combination of features relies on high thresholds and bipolar receptive fields that are (largely) non-overlapping. These properties (high threshold subunits with smaller, non-overlapping receptive fields) are common across multiple species.

Beyond the retina, multiple stages of cascaded nonlinear computation constitutes a ubiquitous motif in the structure and function of neural circuits. The tools we have developed to elucidate hierarchical nonlinear processing in the retina are similarly applicable across neural systems more generally. Thus we hope our work provides mathematical and computational tools for efficiently extracting and analyzing both informative descriptive statistics and hierarchical nonlinear models across many different sensory modalities, brain regions, and stimulus ensembles, thereby furthering our understanding of general principles underlying nonlinear neural computation.

## Methods

### Experiments

Experimental data was collected from the tiger salamander retina using a multi-electrode array (Multi-channel systems), as described elsewhere [9]. Isolated ganglion cells were identified using custom spike sorting software. The stimulus used was a 100Hz white noise bars stimulus, where the luminance of each bar was drawn independently from a Gaussian distribution. Spatially, the stimulus spanned approximately 2.8mm on the retina (50 bars at 55.5*μm*/bar). Intracellular recordings were performed as described elsewhere [73]. Off bipolar cells were identified by their flash response, receptive field size, and level in the retina.

### Regularized spike-triggered analysis

To illustrate the benefits of the regularization terms used to fit the LN-LN models, we apply these penalties to perform regularized spike-triggered analysis. We expect both the STA and the STC eigenvectors to be linear combinations of filters in encoding models with multiple pathways (derivation in Appendix S1). Therefore, we expect certain types of structure in said pathways to persist after the linear combination, assuming the number of pathways is small relative to the stimulus dimension. We thus impose this structure directly on the descriptive statistics through a set of penalty functions (regularizers). For example, these penalties, as functions on a candidate descriptive statistic, could encourage smoothness in the spatial and temporal domains, low rank spatiotemporal structure, or sparsity (i.e. encouraging many filter coefficients to be zero). The proximal algorithms framework allows us to easily incorporate these penalties into our analysis. Below, we formulate optimization problems for regularizing the spike-triggered average and covariance which only require access to the un-regularized estimates. This is useful for the situation where working with the full spike-triggered ensemble or raw dataset is prohibitive due to computational space or time constraints.

### Regularized STA

To compute a regularized STA, without explicitly building an encoding model, we can form an optimization problem that directly denoises the STA:

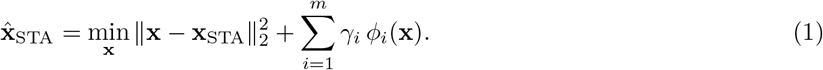

Here, *ϕ_i_(x*) are the regularization penalty functions, with an associated regularization weight *γ_i_*, and x_STA_ is the raw (sample) STA from recorded data (which is noisy due to finite sampling). Note that we use mean squared error to quantify distance from the raw estimate, but other loss functions may be also used. For the penalty functions *ϕ_i_*, we use an *ℓ*_1_ penalty that encourages the estimated filter to be sparse (few non-zero coefficients), and a nuclear norm penalty, which is the sum of the singular values of the spatiotemporal filter **x** when viewed as a spatiotemporal matrix. The nuclear norm penalty is advantageous compared to explicitly forcing the spacetime filter x to be low-rank, as it is a “soft” penalty which allows for many small singular values, whereas explicitly forcing the filter to be low-rank forces those to be zero.

### Regularized STC Analysis

The STC eigenvectors are obtained by an eigendecomposition of the STC matrix **C** [10, 74], which is equivalent to solving an optimization problem:

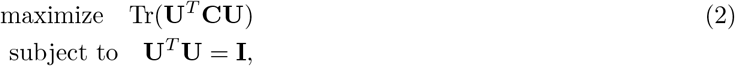

where **U** denotes a matrix whose columns are the orthonormal eigenvectors of **C**. In order to regularize these eigenvectors, we wish to add penalty terms to (2), which precludes a closed form solution to the problem. We circumvent this by reformulating the problem using a convex relaxation. First, we consider the matrix **X = UU**^*T*^, corresponding to the outer product of the eigenvectors. Because of the cyclic property of the trace, namely that Tr(**U**^*T*^**CU**) = Tr(**UU**^T^**C**) = Tr(**XC**), the function to be optimized in (2) depends on the eigenvector matrix **U** only through the combination **X = UU**^*T*^. Thus we can directly optimize over the variable **X**. However, the non-convex equality constraint **U**^*T*^**U** = **I** in (2) is not easily expressible in terms of **X. X** is however a projection operator, obeying **X**^2^ = **X**. We replace this with the constraint that X should be contained within the convex hull of the set of rank-*d* projection matrices. This space of matrices is a convex body known as the *fantope* [75].

The advantage of this formulation is that we obtain a convex optimization problem which can be further augmented with additional functions that penalize the columns of **X** to impose prior knowledge about the structure of the eigenvectors of **C**. Columns of **X** are linear combinations of the eigenvectors of **C**, which are themselves linear combinations of the small set of spatiotemporal filters we are interested in identifying. Therefore, if we expect the spatiotemporal filters of individual biological pathways to have certain structure (for example, smooth, low-rank, or sparse), then we also expect to see those properties in both the eigenvectors and in the columns of **X**.

Putting this logic together, to obtain *regularized* STC eigenvectors, we solve the following convex optimization problem:

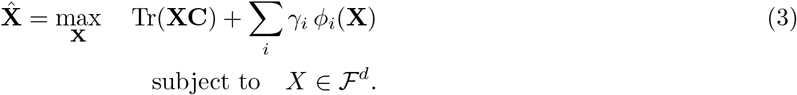

Here **C** is the raw (sample) STC matrix, which is again noisy due to limited recorded data, and 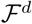 denotes the fantope, or convex hull of all rank *d* projection matrices. Each *ϕ_i_* is a regularization penalty function applied to each of the columns of **X**; i.e. 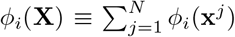 where **x**^*J*^ denotes the *j*’th column of **X.**

Again, we can solve this optimization problem efficiently using proximal consensus algorithms, described below. Common regularization penalties and their corresponding proximal operators are shown in Table S4. The optimization yields a matrix 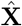 in the fantope 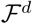, which may itself have rank higher than *d*, so we perform a final eigendecomposition of this matrix to obtain its top-eigenvectors. These eigenvectors constitute our regularized estimate of the eigenvectors of the significant (expansive) STC eigenvectors (to find suppressive directions, one could invert **C** in (3)). A major computational advantage of this formulation is that we only need to store and work with the *N* by *N* raw STC covariance matrix itself, without ever needing access to the spike-triggered ensemble, an *N* by *M* matrix where *M* (the number of spikes) is typically much greater than *N*.

### Proximal operators and algorithms

The framework of *proximal algorithms* allows us to efficiently optimize functions with non-smooth terms. The name *proximal* comes from the fact that these algorithms utilize the *proximal operator* (defined below) as subroutines or steps in the optimization algorithm. For brevity, we skip the derivation of these algorithms, instead referring the reader to the more thorough treatment by Parikh and Boyd [48] or Polson et al. [49]. The proximal operator for a function *ϕ* given a starting point *v* is defined as:

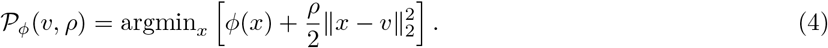

The proximal operator is a mapping from a starting point *υ* to a new point *x* that tries to minimize the function *ϕ(x*) (first term above) but stays close to the starting point *υ* (second term). The proximal operator is a building block that we will use to create more complicated algorithms. We will take advantage of the fact that for many functions *ϕ* of interest to us, we can analytically compute their proximal operators, thus making these operators a computationally cheap building block.

We used these building blocks to solve optimization problems involving the sum of a number of simpler terms:

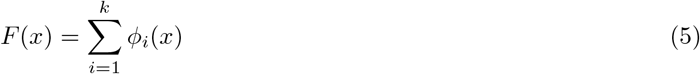

where in our application the *ϕ_i_*’s represent either data fitting terms (e.g. a log-likelihood) or different regularization terms or penalty functions on the parameters, *x*. With respect to learning the parameters of a linear filter in an LN model, the objective consists of a log-likelihood *f*(*x*) along with regularization penalties that impose prior beliefs about the filter, *x*. We focus on two main penalties. Sparsity, which encodes the belief that many filter coefficients are zero, is penalized by the *ℓ*_1_-norm (*ϕ_i_*(*x*) = ||*x*||_1_). Additionally, spatiotemporal filters are often approximately space-time separable (they are well modeled as the outer product of a few spatial and temporal factors). We encoded this penalty by the nuclear norm, *ℓ*_*_, which encourages the parameters *x*, when reshaped to form a spatiotemporal matrix, to be a low-rank matrix (the nuclear norm *ℓ_*_* of a matrix is simply the sum of its singular values). Another natural penalty would be one that encourages the parameters to be smooth in space and/or time, which could be accomplished by applying an *ℓ*_1_ or *ℓ_2_* penalty to the spatial or temporal differences in parameters. As shown below, these types of penalties are easy to incorporate into the proximal algorithm framework. Other commonly used regularization penalties, and their corresponding proximal operators, are listed in Table S4.

The proximal consensus algorithm is an iterative algorithm for solving (5) that takes a series of proximal operator steps. It first creates a copy of the variable *x* for each term *ϕ_i_* in the objective. The algorithm proceeds by alternating between taking proximal steps for each function *ϕ_i_* using that variable copy *x_i_*, and then enforcing all of the different variable copies to agree (reach consensus) by averaging them. The algorithm is:

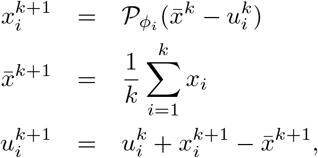

where *i* indexes each of the terms in the objective function, *x_i_* is a copy of the variable, 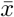 is the average of the variable copies, and *u_i_* is known as a dual variable that can be thought of as keeping a running average of the error between each variable copy and the average. Intuitively, we can think of each variable copy *x_i_* as trying to minimize a single term *ϕ_i_* in the objective, and the average, or consensus 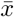 forces the different copies to agree on the best value for the global parameters. After convergence, each copy *x_i_* will be close to the mean value 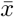, which is the set of parameters that minimizes the original composite objective.

This algorithm has a number of desirable properties. First, the updates for each term *x_i_* can be carried out in parallel, therefore allowing for speedups when run on a cluster or multi-core computer. Second, it converges even when terms in the objective are non-differentiable. Due to the repeated application of the proximal operator, this algorithm works best when the terms *ϕ_i_* have proximal operators that are easy to compute.

This is exactly the case for the regularization terms described above: for the *ℓ*_1_ norm, the proximal operator corresponds to soft thresholding of the parameters. For the nuclear norm, the proximal operator corresponds to soft thresholding of the singular values of parameters reshaped as a matrix. Occasionally, the proximal operator may not have a closed form solution. In this case, the proximal step can be carried out through gradient based optimization of (4) directly. This is the case for some log-likelihoods, such as the log-likelihood of a particular firing rate under Poisson spiking. In this case, gradient step based optimization of (4) often dominates the computational cost of the algorithm. As many methods for fitting neural models involve gradient step updates on the log-likelihood, such methods can then be augmented with additional regularization terms with no appreciable effect on runtime, by using proximal consensus algorithms for optimization. Our code for solving formulating and solving optimization problems using proximal algorithms is provided online at https://github.com/ganguli-lab/proxalgs.

### LN-LN models

In this section, we specify the mathematical formulation of our LN-LN models. The model takes a spatiotemporal stimulus, represented as a vector **x**, and generates a predicted firing rate, *r*(**x**). First, the stimulus is projected onto a number of subunit filters. The number of subunits is a hyper-parameter of the model, chosen through cross validation (we repeatedly fit models with increasing numbers of subunits until held-out performance on a validation set decreases). If we have *k* subunits, then the stimulus is projected onto each of the *k* filters: w_*i*_ for *i* = 1, …, *k*. These projections are then passed through separate subunit nonlinearities. The nonlinearities are parameterized using a set of Gaussian basis functions (or bumps) that tile the relevant input space [12, 20], this enforces the nonlinearities to be smooth. We typically use *p* = 30 evenly spaced Gaussian bumps that tile the range spanned by the projection of the stimulus onto the linear filter (results were not sensitive to the number of bumps over a range of 10–30 bumps). For example, a nonlinearity *h*(*u*) is parameterized as

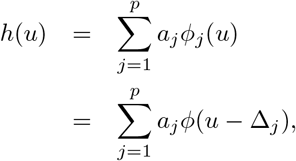

where *ϕ* is the basis function, e.g. *ϕ(x*) = exp(–*x*^2^), Δ_*j*_ indicates the spacing between the basis functions, *p* is the number of bases used, and *a_j_* is a weight on that particular basis function. Since the basis functions and spacings are fixed beforehand, the only free parameters are the *α_j_*’s. For subunit *i*, the corresponding nonlinearity has a set of weights *a_i,j_* for *j* = 1,…, *p*.

The output of the *k* subunits is then summed and passed through a final nonlinearity. This final nonlinearity is parameterized as a soft rectifying function *r(x*) = *g* log(1 + *e^x-θ^*) with two parameters: *g* is an overall gain and *θ* is the threshold. The full LN-LN model is then given by:

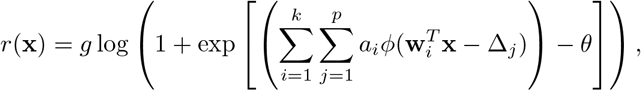

where the parameters to optimize are the subunit filters w_*i*_ for *i* = 1,…, *k*, subunit nonlinearity weights *a_i,j_* = for *j* = 1, …, *p*, and final nonlinearity parameters *θ* and *g*.

We optimize the parameters using a maximum likelihood objective assuming a Poisson noise model for spiking. Rather than optimize all of the parameters simultaneously, we alternate between optimizing blocks of parameters (joint optimization using gradient descent was prone to getting stuck at solutions that were less accurate). That is, we alternate between optimizing three blocks of parameters: the subunit filters w_*i*_, the subunit nonlinearities *a_ij_*, and the final nonlinearity parameterized by *θ* and *g*. We optimize each block of parameters by minimizing the negative log-likelihood of the data plus any regularization terms using proximal algorithms. The subunit filters are the only parameters with regularization penalties (the nuclear norm applied to the filter reshaped as a spatiotemporal matrix and the *ℓ*_1_ norm), to encourage space-time separability and sparseness of the filters. The proximal operator for each of these regularization penalties is given in Table S4, and the proximal operator for the log-likelihood term (which does not have a closed-form solution) is solved using gradient descent. In addition, after optimizing the block of parameters corresponding to the subunit filters, we rescale them to have unit norm before continuing the alternating minimization scheme. This ensures that the distribution of input to the nonlinearities spans the same range, and gets rid of an ambiguity between the scale of the subunit filters and the scale of the domain of the subunit nonlinearity. We find that the parameters converge after several rounds of alternating minimization, and are robust with respect to random initialization of the parameters.

### Subspace overlap

We quantify the overlap between two subspaces as the average of the cosine of the principal (or canonical) angles between the subspaces. The principal angles between two subspaces 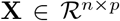 and 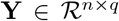 generalize the idea of angles between vectors. Here we describe a pair of *p* and *q* dimensional subspaces in *n* dimensional space as the span of the columns of the matrices **X** and **Y**. Assuming without loss of generality that *p* ≤ *q*, then we have *p* principal angles *θ*_1_,…, *θ_p_* that are defined recursively for *k* = 1,…,*p* as:

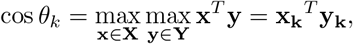

subject to the constraints that the vectors are unit vectors (x^*T*^ × = y^*T*^y = 1) and are orthogonal to the previously identified vectors 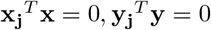 for *j* = 1, 2,…, *k* – 1). That is, the first principal angle is found by indentifying a unit vector within each subspace such that the correlation, or dot product, between these vectors (these are known as the principal vectors) is maximized. This principal angle is then the inverse cosine of the dot product. Each subsequent principal angle is found by performing the same maximization but restricting each new pair of vectors to be orthogonal to the previous principal vectors in each subspace. The principal angles can be efficiently computed via the QR decomposition [76]. We define subspace overlap as the average of the cosine of the principal angles, 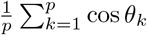 This quantity is at most 1 (for two subspaces that span the same space), and at least 0 (for two orthogonal subspaces that share no common directions).

## Acknowledgments

The authors would like to thank Ben Naecker, Ben Poole, and Lane McIntosh for discussions as well as Stéphane Deny, Ben Naecker, and Alex Williams for comments on the manuscript.

## S1 Appendix Relationship between descriptive statistics and encoding models

Here, we derive the relationship between the pathways of any differentiable encoding model and spike-triggered statistics under Gaussian noise stimulation. We represent a visual stimulus as an *N* dimensional vector **x**. We view a functional neural model as an arbitrary nonlinear function *r* = *f*(x), over *N* dimensional stimulus space, where *r* determines the probability that the neuron fires in a small time window following a stimulus x: *r*(x) = *p*(spike | x). The derivation will show how the STA is related to the *gradient* of the model ∇*r*(x), and the STC is related to the *Hessian*, ∇^2^*r*(x).

The STA and STC are the mean and covariance, respectively, of the spike-triggered stimulus ensemble, which reflects the collection of stimuli preceding each spike [3]. This distribution over stimuli, conditioned on a spike occurring, can be expressed via Bayes rule,

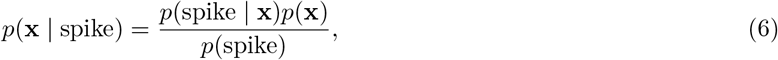

where *p*(x) is the prior distribution over stimuli and *p*(spike) is the average firing probability over all stimuli. Here, we assume a white noise stimulus distribution, in which each component of x is chosen independently from a Gaussian distribution with zero mean and unit variance. The STA and STC are given by

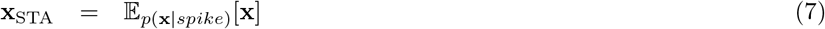

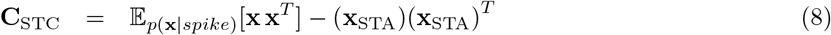

Focusing first on the STA:

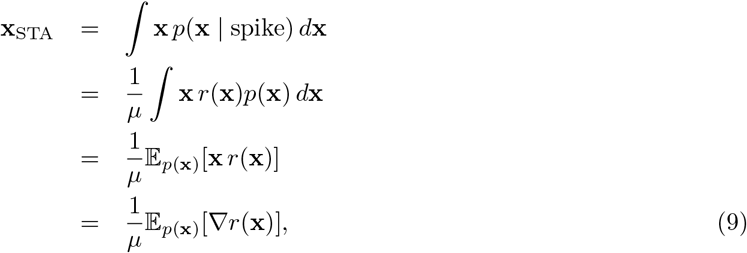

where *μ* = *p*(spike) is the overall probability of spiking. The last step in the derivation uses Stein’s lemma, which states that 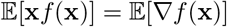 if the expectation is taken over a multivariate Gaussian distribution with identity covariance matrix, corresponding to our white noise stimulus assumption. This calculation thus yields the simple statement that the spike-triggered average is proportional to the gradient (or gain) of the response function, averaged over the input distribution [77]. Applying Stein’s lemma again yields an expression for the STC matrix:

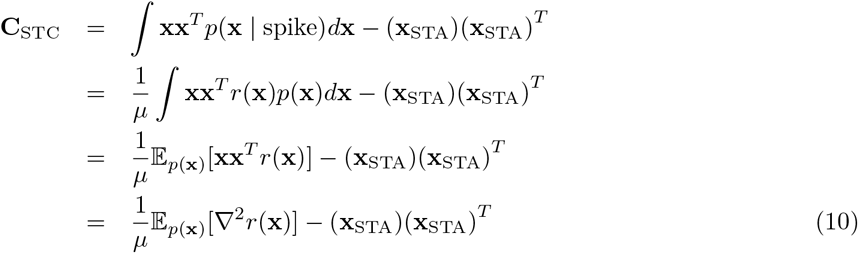

Intuitively, these results state that the STA is related to the slope (first derivative) and the STC is related to the Hessian curvature (matrix of second derivatives) of the multi-dimensional nonlinear response function *r*(x).

For example, consider a linear-nonlinear model *r* = *f*(**w**^*T*^**x**) which has the following gradient: ∇*r*(x) = *f*′(**w**^*T*^**x**)**w** and Hessian: ∇^2^*r*(**x**) = *f*″(**w**^*T*^**x**)**ww**^*T*^. Plugging these expressions into equations (9) and (10) reveals that the STA is proportional to **w** and the STC is proportional to **ww**^*T*^. Therefore, we recover the known result [2] that the STA of the LN model is proportional to the linear filter, and there will be one significant direction in the STC, which is also proportional to the linear filter (with mild assumptions on the nonlinearity, *f*, to ensure that slope and curvature terms in (9) and (10) are non-zero).

We can extend this to the case of a multilayered circuit with *k* pathways, each of which first filters the stimulus with a filter **w**_1_…**w**_*k*_. Regardless of how these pathways are then combined, we can write this circuit computation as *r* = *f*(**W**^*T*^**x**) where **W** is a matrix whose columns are the *k* pathway filters, and *f* is a *k*-dimensional time-independent (static) nonlinear function. We can think of the *k* dimensional vector **u** = **W**^*T*^**x** as the activity pattern across each of the *k* pathways before any nonlinearity. The gradient for such a model is ∇*r*(**x**) = **W**^*T*^**∇f(u)**, where **∇f(u)** is the gradient of the *k*-dimensional nonlinearity. Using equation (9), the STA is then a linear combination of the pathway filters:

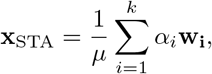

where the weights are given by

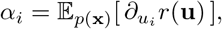

and correspond to the average sensitivity, or slope of the neural response *r* with respect to changes in the activity of the *i^th^* filter.

The Hessian for the multilayered model is **∇**^2^*r*(**x**) = **W∇**^2^**fW**^*T*^, where **V**^2^**f** is the *k*-by-*k* matrix of second derivatives of the *k*-dimensional nonlinearity *f*(**u**). From equation (10), the STC is then given by:

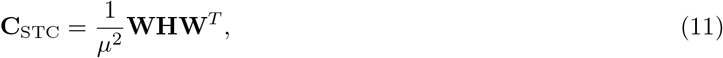

where the *k*-by-*k* matrix **H** is:

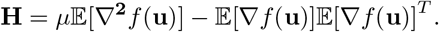

This expression implies that nontrivial directions in the column space of C_STC_ correspond to (span the same space as) the column space of **W**. Therefore, the significant eigenvectors of the STC matrix will be linear combinations of the *k* pathway filters, and the number of significant eigenvectors is at most *k*.

Note that equations (9) and (10) are valid for *any* differentiable model, including those with more than two layers, divisive interactions, feedback, and so on.

## S2 Figure Example bipolar cell receptive fields

**Fig 11.**
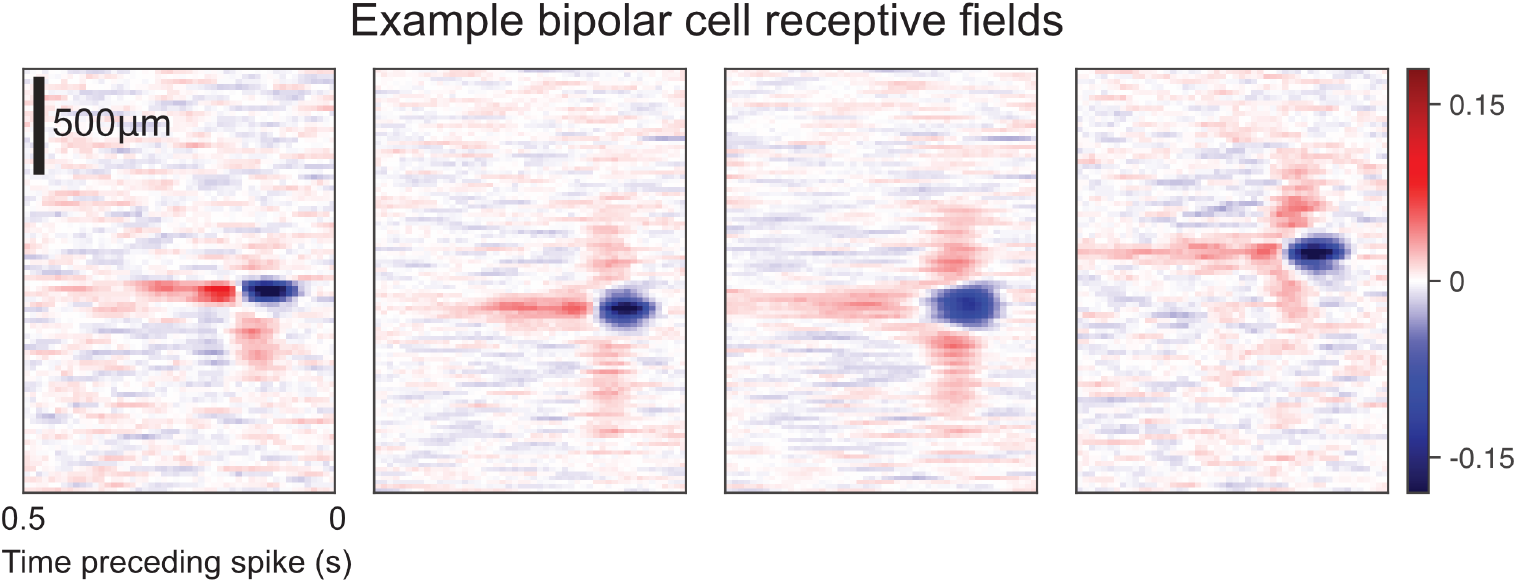
Example bipolar cell receptive fields. Each panel shows a spatiotemporal receptive field of a bipolar cell, recorded intracellularly from the salamander retina (see Methods).

## S3 Figure Subunit and retinal ganglion cell types

**Fig 12.**
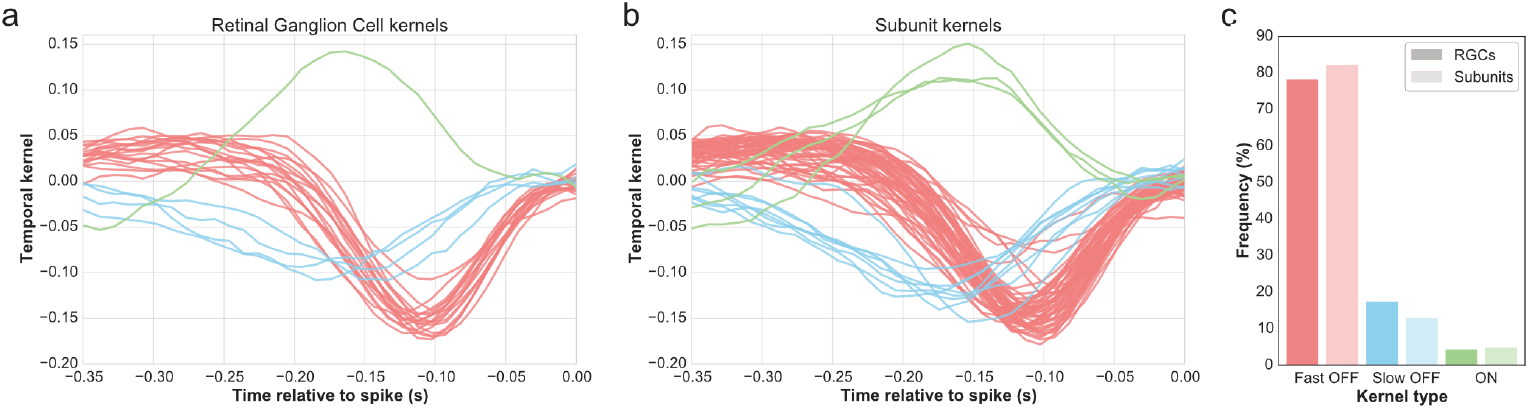
Cell type classification for salamander ganglion cell and subunit filters. (a) K-Means clustering applied to the temporal kernel (temporal component of the spatiotemporal receptive field) of *n* = 23 recorded retinal ganglion cells. (b) K-Means clustering applied to temporal kernels of *n* = 92 model-identified subunits. (c) Frequency of the different cell types, both for RGCs and subunits.

## S4 Table Proximal operators for common regularization penalties

**Table 1.**
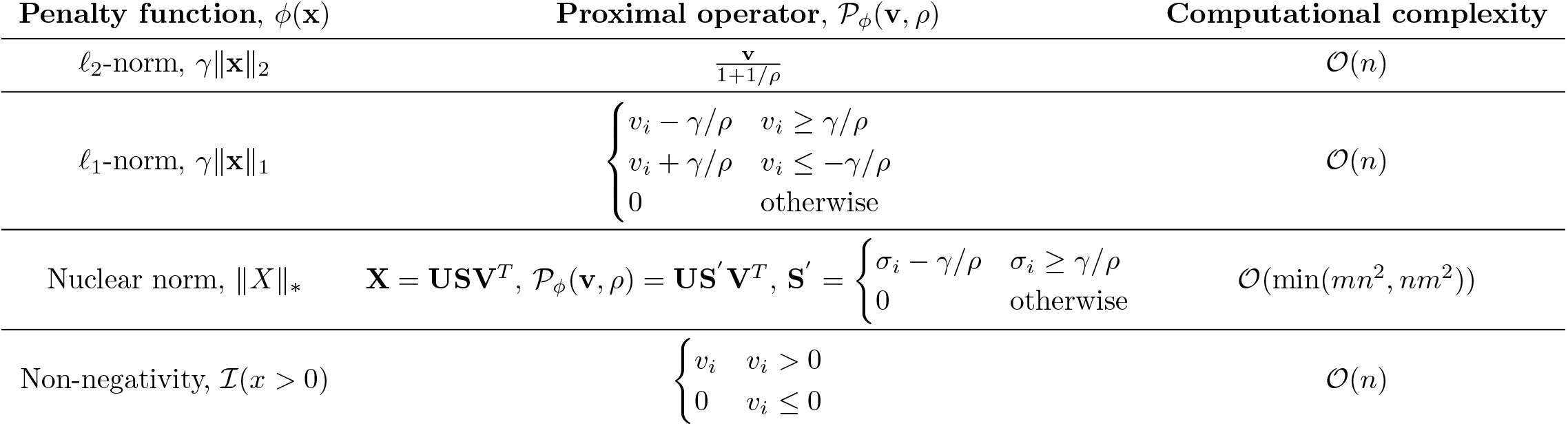
Common regularization penalties and their proximal operators (in closed form).

## Footnotes

1 Note that subunits that combined *linearly* would be indistinguishable from a computational perspective. Due to the roughly linear integration [9] that occurs at bipolar cells, we (computationally) distill mechanisms in photoreceptors and inhibitory horizontal cells into a single spatiotemporal filter with positive and negative elements that gives rise to bipolar cell signals.

